# Acetylation of WCC is dispensable for the core circadian clock but differentially regulates acute light responses in *Neurospora*

**DOI:** 10.1101/2023.11.29.569266

**Authors:** Bin Wang, Mark E. Edamo, Xiaoying Zhou, Ziyan Wang, Scott A. Gerber, Arminja N. Kettenbach, Jay C. Dunlap

## Abstract

In the *Neurospora* circadian system, the White Collar Complex (WCC) formed by WC-1 and WC-2 drives expression of the *frequency* (*frq*) gene whose product FRQ feedbacks to inhibit transcriptional activity of WCC. Phosphorylation of WCC has been extensively studied, but the extent and significance of other post-translational modifications (PTM) has been poorly studied. To this end, we used mass-spectrometry to study alkylation sites on WCC, resulting in discovery of nine acetylation sites. Mutagenesis analysis showed most of the acetylation events individually do not play important roles in period determination. Moreover, mutating all the lysines falling in either half of WC-1 or all the lysine residues in WC-2 to arginines did not abolish circadian rhythms. In addition, we also found nine mono-methylation sites on WC-1, but like acetylation, individual ablation of most of the mono-methylation events did not result in a significant period change. Taken together, the data here suggest that acetylation or mono-methylation on WCC is not a determinant of the pace of the circadian feedback loop. The finding is consistent with a model in which repression of WCC’s circadian activity is controlled mainly by phosphorylation. Interestingly, light-induced expression of some light-responsive genes has been modulated in certain *wc-1* acetylation mutants, suggesting that WC-1 acetylation events differentially regulate light responses.

## Introduction

Circadian rhythms that have been found in most eukaryotes and certain prokaryotes orchestrate a wide variety of behavioral, metabolic, and molecular processes (1), allowing organisms to effectively utilize environmental resources. Dysfunction of the endogenous clock in humans has been implicated in many health issues and even severe disorders (2). Mechanistically, circadian clocks are composed of interlocked positive and negative element complexes: The latter progressively inhibit their own expression through deactivating the transcriptional activity of the former. *Neurospora crassa*, a genetically trackable model fungus sharing many basic cellular features with mammals, has been employed for circadian research for decades, facilitating great contributions to our understanding of the core clock operation (3–5). In *Neurospora*, the White Collar Complex (WCC) assembled from WC-1 and WC-2 acts as a signaling pivot for synchronizing the core oscillator that operates in darkness and can respond to environmental light signals (6). There exist two independent DNA elements in the promoter of the central circadian pacemaker gene *frequency* (*frq*): In constant darkness, WCC drives circadian expression of *frq* by binding to the *Clock box* (*C-box*) DNA element (7), while it associates with the *Proximal Light-Response Element* (*pLRE*) to activate *frq* transcription upon light exposure (8, 9).

FREQUENCY (FRQ), the gene product of *frq*, nucleates formation of a multi-component complex, FFC (FRQ-FRH complex), with FRH (FRQ-interacting RNA helicase) (10–12) and CKI (Casein Kinase I) (13). In circadian cycles, FFC promotes phosphorylation of WCC at over 95 residues, most of which are involved only in downregulation of circadian amplitudes (expression levels of circadian genes) but not in determining period length (14). However, a circadian repression in the core oscillator arises from phosphorylation of a small group of these residues (13–16). FRQ itself also undergoes successive and extensive phosphorylations at over 110 residues (17, 18), many of which markedly impact the pace of the core circadian oscillator including prolonging or shortening the period length to variable extents as has been established by mutagenesis analyses of *frq* (19–21).

The mammalian functional orthologs of WC-1 and WC-2 are Bmal1 and its interacting partner, CLOCK (transcription factors bound to the *E-box* DNA element), that drive expression of circadian negative-element proteins Periods (Pers) and Cryptochromes (Crys), all of which feedback to repress transcriptional activity of the Bmal1/CLOCK complex (reviewed in [22]). In mammals, multiple types of post-translational modifications (PTMs) have been identified on Bmal1 and CLOCK, including SUMOylation, ADP-ribosylation, O-GlcNAcylation, acetylation, ubiquitination, S-nitrosylation, and phosphorylation (23–28). However, which PTM plays the major role in modulating the pace of the circadian oscillator remains unclear, probably partially due to gene redundancy of the circadian components. It was proposed that in the repression phase of the clock, CLOCK-mediated acetylation of Bmal1 at K537 in mouse (equivalent to K538 of human Bmal1) is required for Cry-mediated repression of Bmal1/ CLOCK (23). But this conclusion has been challenged by several lines of evidence: 1) Acetylation of Bmal1 occurs during the activation phase of a circadian cycle (29, 30); 2) Acetylation of K537 is mediated by Tip60 instead of CLOCK (24); 3) The K537R mutant of Bmal1 that abrogates K537 acetylation behaves like partial loss-of-function rather than being more active than WT (23), as would be expected for loss of a negative-acting PTM. In *Neurospora*, although phosphorylation has been found to be a determinant in depressing transcriptional activity of WC-1/ WC-2 (see above), acetylation has also been detected in biochemical assays (31). Also, physical association between WC-1 and the histone acetyl-transferase NGF-1 has been reported as required for light responses (30), and has been associated with light-induction of gene expression. Despite these hints however, the exact acetyl-sites are wholly unknown let alone their possible roles in the feedback mechanism dictating the core clock.

To explore possible alternative regulatory mechanisms in the circadian feedback loop we studied WCC using mass-spectrometry to identify PTMs other than phosphorylation and found multiple acetylation and mono-methylation sites. Surprisingly however, clock functions including period length and phase were little affected in most *wc-1 or wc-2* mutants bearing single mutations at the acetylated or mono-methylated residues. Furthermore, even when all lysines on WC-1 or WC-2 were covered in the mutagenesis analysis, robust circadian oscillations were still observed. Altogether, the data here suggest that acetylation does not play important roles in modulating circadian periods of *Neurospora*, and acetylation of the circadian positive element proteins may be not a conserved mechanism for tuning their circadian activities.

## Results

### Identification of acetylated residues on WC-1 and WC-2

In order to probe post-translational modifications (PTMs) beyond phosphorylation on the White Collar Complex (WCC), WC-1 and WC-2 were purified as from cultures grown in constant light (LL) and analyzed by mass spectrometry. Spectra were searched for evidence of acetylation, (mono-, di-, and tri-) methylation, and ubiquitination, all of which have been reported on Bmal1/CLOCK (see Introduction), the mammalian functional orthologs of *Neurospora* WC-1/WC-2. Interestingly, six acetylation sites were found on WC-1 and three on WC-2 (Fig. 1A, Supporting Figure 2, and Supporting Table S1) with the coverage of 99.1% for WC-1 and 97.7% for WC-2 (Supporting Fig. 2). This result is consistent with previous literature that used Western blotting to show that WC-1 is an acetylated protein (31). Mono-methylation was also detected on WCC (Supporting Table S1 and see below); however, we did not retrieve any ubiquitinated peptides from WC-1 or WC-2, probably in part because we did not attempt to inhibit proteasomes in the cells from which WCC was isolated. The data here indicate that the positive-element complexes in *Neurospora* and mammals (see Introduction) are both alkylated at multiple residues.

**Figure 1.**
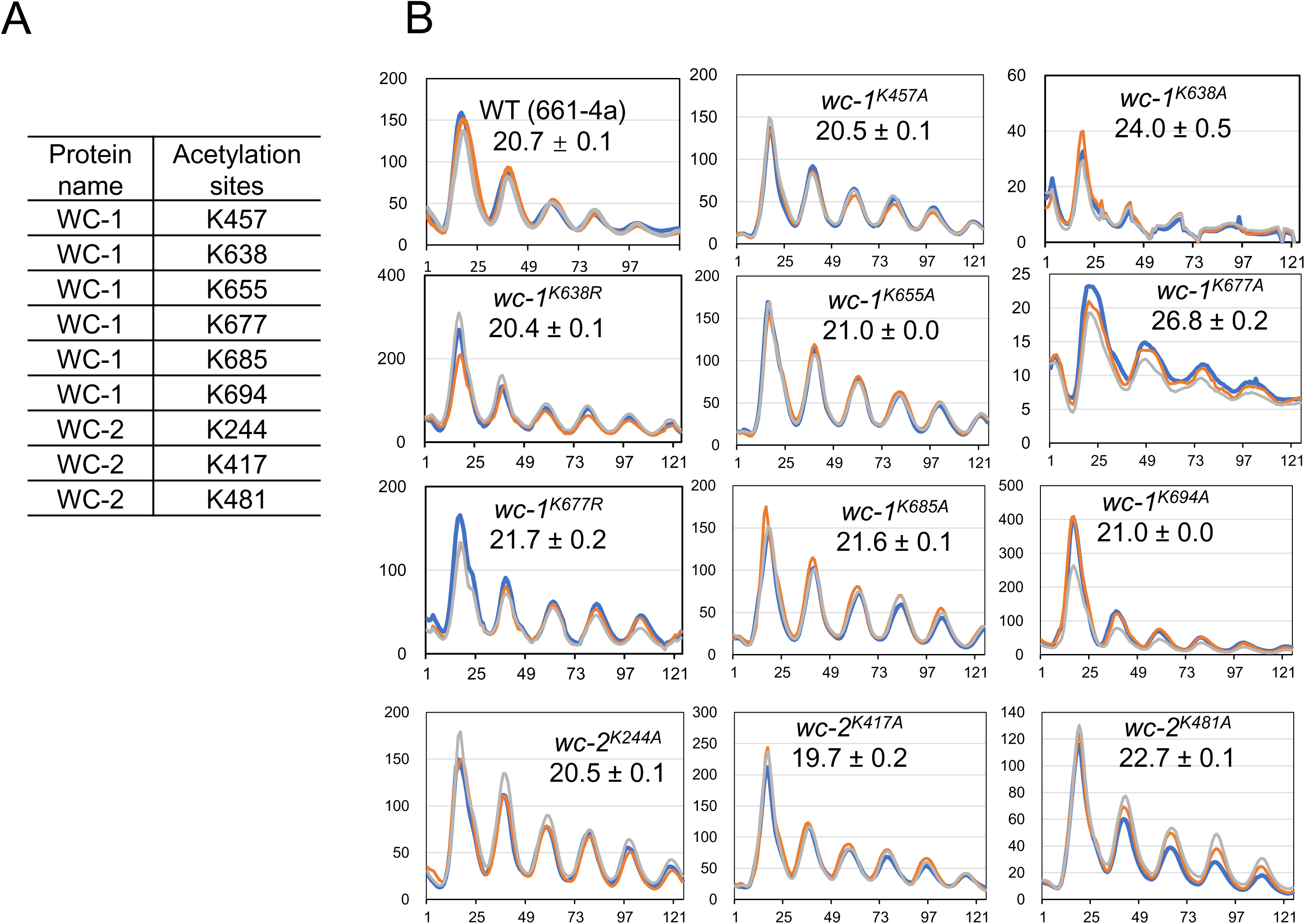
Identification and mutagenesis study of acetylation on WC-1 and WC-2. (**A**) List of identified acetylation sites on WC-1 and WC-2 by mass spectrometry (**B**) Luciferase analysis of *wc-1* mutants bearing alanine mutants to the mono-methylation sites as stated. Strains were grown at 25 °C plus light overnight, and bioluminescence signals were recorded hourly with a CCD (charge-coupled device) camera after transfer of the strains to the dark also at 25 °C. Three replicates (in different colors) were plotted with the x-axis and y-axis meaning time (in hours) and arbitrary units of the bioluminescence-signal intensity, respectively. Period length was calculated from three replicates and reported as the average +/- the standard error of the mean (SEM). All *wc-1* or *wc-2* mutants throughout this study were targeted to their native loci with a V5 at the C-terminus of WC-1 while WC-2 untagged.

### Mutagenesis analysis of acetylation sites on WCC

To examine possible role(s) of WCC acetylation in the circadian clock, mutagenesis analysis (Fig. 1B) was performed to eliminate acetyl-sites found on WC-1 and WC-2 (Fig. 1A). Mutations in most of the WC-1 or WC-2 acetyl-sites, when changed individually (to alanine,) did not significantly impact period length (Fig. 1B) except for K677 on WC-1. *wc-1^K677A^* showed a period of 26.8 hrs with a low amplitude (compare y-axes), long period and low amplitude being characteristic of *wc-1* or *wc-2* mutants with partially compromised transcriptional activity. For example, ER24, a *wc-2* mutant bearing a point mutation in the Zinc-finger DNA-binding domain rendering it partially defective in DNA binding, exhibits a markedly long period (28.6 hrs) (32); several *wc-1* mutants having partial deletions in the coactivator-recruitment domain show an approximately 25-hrs period (33). On the contrary, *wc-1* or *wc-2* mutants with enhanced circadian activity always display shortened periods with a high amplitude (14). K677 is situated in the linker region between PASB and PASC, both of which are required for WC-1 to be functional in driving *frq* expression and WC-2 interaction (34). Therefore, *wc-1^K677A^* seems to be a partial loss-of-function mutant based on the long period as well as the low amplitude. Another piece of evidence comes from substitutions of arginine for lysine, a change that preserves the positive charge but prevents acetylation. *wc-1^580-1167K0R^*, in which all lysines (including K677) falling in aa 580-1167 of WC-1 were mutated to arginines, did not show a period-length adjustment (see below). Overall, the mutagenesis data suggests that most single acetylation events identified on WCC do not significantly impact the pace of the core oscillator.

### Biochemical analysis of acetylation on WCC

To follow the acetylation status of WC-1 and WC-2 more conveniently, we performed Western blotting (WB) with several commercially available antibodies. The three antibodies that we tried have been highly rated or cited by users for successfully detecting lysine acetylation on proteins from a broad range of organisms (see Experimental Procedures). To prevent false positive detection in Western blotting from nonspecific binding, WC-1 (tagged with 3xFLAG at the C terminus) was first isolated from the lysate by immunoprecipitation (IP) prior to Western blotting with acetylated lysine-specific antibodies. The assay included three controls: WT (untagged) served as a negative control for Western blotting and IP, a positive control for acetyl-lysine-specific antibodies came from a *Neurospora* strain ectopically expressing a second copy of histone H4 with a C-terminal 3xFLAG at the *csr-1* locus, and another strain as the negative control for acetylation bore a second copy of histone H4 (also with a C-terminal 3xFLAG and at *csr-1*) carrying K5A, K8A, K12A, and K16A mutations, four acetylation sites conserved widely among diverse organisms (35, 36). Following FLAG IP and Western blotting, specific bands with expected sizes corresponding to histone H4, histone H4^K5A, K8A, K12A, K16A^, and WC-1 appeared in samples from the transgenic strains, which were not seen in the untagged strain (Fig. 2A, the FLAG blot). Strong and specific histone H4 bands were observed with each of the three acetylated lysine (AcK)-specific antibodies (Fig. 2A, AcK antibodies from Cell Signaling Technology [CST], Santa Cruz Biotechnology [San Cruz], or Invitrogen). At the same position of histone H4, only very faint bands were seen from the sample of histone H4^K5A, K8A, K12A, K16A^ (denoted by the lower red box in Fig. 2A), probably due to weak non-specific reaction between the highly abundant unacetylated (K->A) histone H4 and the AcK antibodies. However, on any of the three Ack blots, no acetylation signal was detected at the position where WC-1 is supposed to appear (defined by the upper red box across the FLAG and three AcK blots in Fig. 2A). The strong and specific signals from the positive control (3xFLAG-tagged histone H4) indicate that the three commercial antibodies against acetylated lysines worked nicely for proteins expressed and extracted from *Neurospora* but also suggests that acetylated WC-1 detected by mass spectrometry (Fig. 1A and Supporting Figure 1) may only represent a small fraction of all the WC-1 molecules, consistent with the absence of circadian period changes in strains bearing single mutations to the acetylation sites of WC-1 or WC-2 (Fig. 1B). If control of WCC’s circadian activity involves acetylation, its level should oscillate in a circadian cycle, like what was seen for WCC phosphorylation (represented by WC-2 phosphorylation) (14, 15). To test this possibility, V5-tagged WC-1 was pulled down with V5 resin and then blotted with the three AcK antibodies that have been validated (Fig. 2A). The strong and specific WC-1^V5^ and WC-2 bands were noticed after IP-enrichment but acetylation signals were not detected at the positions of WC-1 and WC-2 on any of the three Ack blots at any examined circadian time points even at DD24 when WCC is fully repressed (Fig. 2B). If acetylation of WCC is a driving mechanism for the in the circadian feedback loop, FRQ as the central negative-element protein for the core oscillator might control acetylation of WCC as it does phosphorylation (14). To this end, FRQ was overexpressed by an inducible promoter, *qa-2*, in the dark for 16 hrs, and WC-1 (V5 tagged) along with WC-2 was isolated (by V5 IP) and blotted for the AcK antibodies. Relative to the no QA sample, the level of WC-1 and WC-2 was markedly and characteristically enhanced by quinic acid (QA)-induced FRQ overexpression, which is consistent totally with previous literature (14, 37) showing that FRQ promotes WCC phosphorylation, makes it leave the chromatin, and thereby protects it from being degraded in transcription, leading to its accumulation. Consistent with the data in Figs. 2A and 2B, WCC acetylation remains undetectable even when FRQ was overexpressed (Fig. 2C). Collectively, the biochemical data here suggest that acetylated WCC detected by mass spectrometry may only account for a very small fraction of the WCC population, and WCC activity seems unlikely to be determined by acetylation due to its low penetrance.

**Figure 2.**
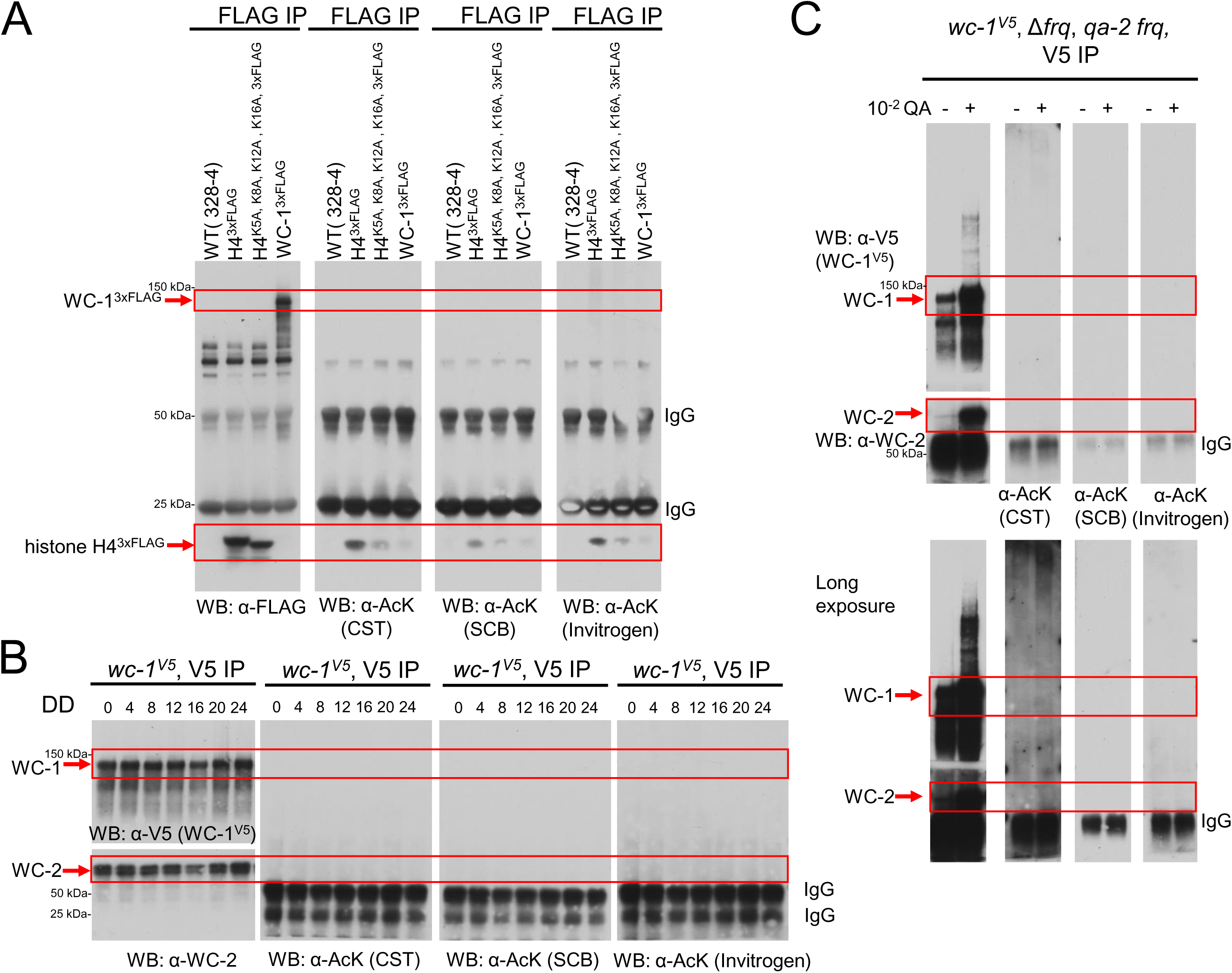
Acetylation of WC-1 and WC-2 was not detected by Western blotting. (**A**) Quality control for acetyl-lysine antibodies. WT (328-4, untagged) served as a negative control for WB and IP, a positive control for acetyl-lysine-specific antibodies came from a *Neurospora* strain ectopically expressing a second copy of histone H4 with a C-terminal 3xFLAG at the *csr-1* locus, and another negative control for acetylation is a strain that bore a second copy of histone H4 (also with a C-terminal 3xFLAG and at *csr-1*) carrying K5A, K8A, K12A, and K16A mutations, four acetylation sites conserved widely among diverse organisms. WC-1 (tagged with 3xFLAG at the C terminus) was isolated first by immunoprecipitation (IP) with FLAG resin. The three commercial acetyl-lysine antibodies were purchased from Cell Signaling Technology [CST], Santa Cruz Biotechnology [SCB], and Invitrogen and used in Western blotting at a dilution of 1:1,000. All *Neurospora* samples were cultured at 25 °C with light for approximately 24 hrs. (**B**) Acetylation of WC-1 and WC-2 was not detected by Western blotting from samples collected in a 24-hr time course (with an interval of 4 hrs). V5-tagged WC-1 (along with WC-2) was isolated from the cell by V5 immunoprecipitation. Western blotting was performed with indicated antibodies. (**C**) Overexpression of FRQ does not induce WCC acetylation. In *wc-1^V5^*, Δ*frq*, *qa-2 frq*, FRQ was overexpressed by adding QA to the 0.1% glucose LCM medium at the final concentration of 10^-2^ M and immediately transferring the culture to the dark at 25 °C for 16 hrs. The same strain cultured under the identical condition but without QA throughout served as the negative control. WC-1 (V5 tagged) was isolated by immunoprecipitation with V5 resin and then Western blotting was carried out with indicated antibodies.

### Phenotypes of *wc-1* and *wc-2* mutants covering all lysine residues

To test combinatorial effects of acetylation events, if any, and also to study WC-1 acetylation events that might be missed in mass spectrometry, *wc-1^K0R^*in which all lysines (a total of 49) of WC-1 were mutated to arginines was generated and assayed. *wc-1^K0R^* displayed an arrhythmic clock in the luciferase assay which monitors transcription activity of WCC at the *frq* promoter (Fig. 3A left), and levels of WC-1, WC-2, and FRQ in *wc-1^K0R^* were all comparable to those in WT (Fig. 3A right). The arrhythmicity and reduced signal intensity (comparing y-axes) in *wc-1^K0R^* are likely due to the large number of mutations (49 in total) that were introduced in the protein, reminiscent of what was seen in another mutant of *wc-1*, *wc-1^113A^*, in which all the 80 phosphosites along with 33 additional Ser or Thr were mutated together to alanines (14).

**Figure 3.**
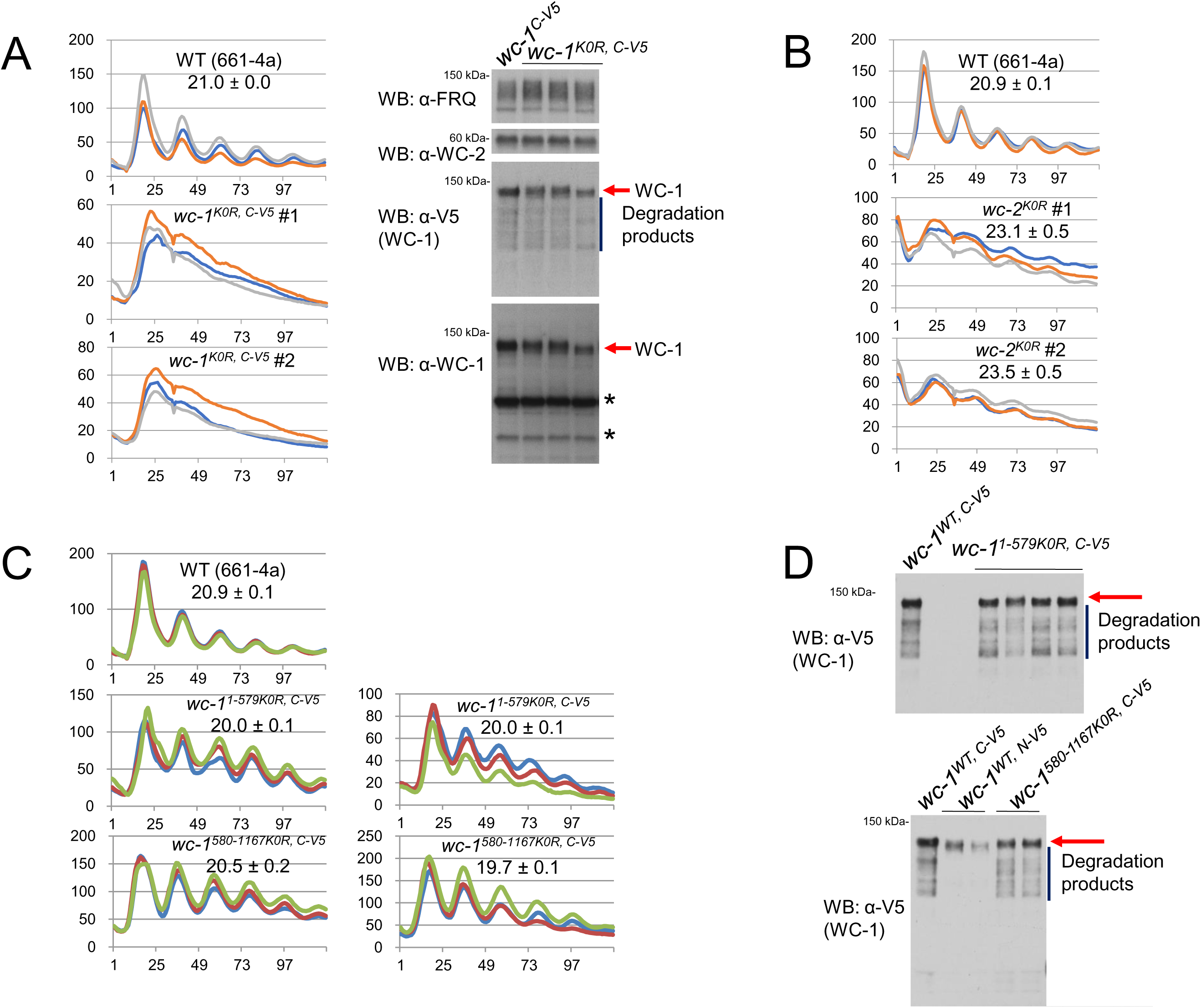
Luciferase analysis of *wc-1^K0R^* and *wc-2^K0R^*. “K0R” means that all lysine residues (in WC-1 or WC-2) were mutated to arginines. Acetylation of WC-1 and WC-2 was not detected by Western blotting. (**A**) Left, luciferase analysis of *wc-1^K0R^*; right, expression of FRQ and WC-1 in *wc-1^K0R^*. (**B**) luciferase analysis of *wc-2^K0R^*. Clock synchronization was done by growing cultures at 25 °C with light overnight (∼ 16 hrs), and bioluminescence signals were recorded hourly with a CCD camera after moving the strains to darkness (at 25 °C). Three replicates were plotted with the x-axis and y-axis representing hours and arbitrary signal intensities, respectively. Period length was computed from the three replicates and presented as the average +/- the standard error of the mean (SEM). WC-1 in *wc-1^K0R^*bear a V5 tag at the C-terminus while WC-2 in *wc-2^K0R^* is tagless. *wc-1^1-579K0R^* and *wc-1^580-1167K0R^* show a wild-type period length and normal WC-1 expression. (**C**) Luciferase analysis of *wc-1^1-579K0R^* and *wc-1^580-1167K0R^* at 25 °C. Synchronization, signal recording, and data plotting follow the same way as figures above. Three replicates (in three colors) were plotted with the x-axis and y-axis meaning time (in hours) and arbitrary light-signal intensities, respectively. (**D**) Expression of WC-1 in *wc-1^1-579K0R^* (upper) and *wc-1^580-1167K0R^* (bottom), both of which have a V5 at the C-terminus. *wc-1^V5^*, the control, also has a C-terminal V5 tag. Protein was extracted from cultures grown in the light at 25 °C. Red arrows point to the full-length WC-1 bands, and diffused bands (denoted with a vertical line) are degradation products of WC-1^V5^.

To examine the role of WC-2 acetylation in the core clock operation, we engineered *wc-2^K0R^* in which all the lysine residues (25 in total) of WC-2 regardless of their acetylation status were mutated to arginines. *wc-2^K0R^* displayed circadian rhythms for four days though its period length was lengthened slightly by ∼ two hrs (Fig. 3B) and the waveform of the rhythm was severely affected. While this likely reflects a side effect of the large number of mutations in the protein, the data indicate that WC-2 acetylation is not required for rhythmicity.

### Mutation of all lysines in either half of WC-1 did not alter period length

To restore circadian oscillations in *wc-1^K0R^* that was arrhythmic in the luciferase assay (Fig. 3A), two additional *wc-1* mutants derived from *wc-1^K0R^* were engineered: *wc-1^1-579K0R^*bearing all lysines from aa 1-579 (16 lysines) mutated to arginine while keeping aa 580-1167 unchanged and *wc-1^580-1167K0R^* in which aa 1-579 remains WT but all lysines located in aa 580-1167 (33 lysines) were mutated to arginines. Both *wc-1* mutants, *wc-1^1-579K0R^* and *wc-1^580-1167K0R^*, showed a robustly oscillating clock with marginally shortened period length (Fig. 3C). The level of WC-1 in *wc-1^1-579K0R^*and *wc-1^580-1167K0R^* is almost identical to that in WT (Fig. 3D). Taken together, these data do not indicate an essential role for acetylation in modulating the circadian activity of WC-1 nor in controlling the operation of the core oscillator in *Neurospora*.

### Acetylation of WCC modulates expression of certain WCC target genes upon light exposure

WCC is not only the central transcriptional activator for the clock but also directly and strongly induces transcription of light responsive genes genome-wide upon a light pulse (38). To test whether expression of light-inducible genes is affected by acetylation events on WC-1 or WC-2 (Fig. 1), the acetylation mutants of *wc-1* and *wc-2* were cultured in the dark for ∼ 24 hrs and thereafter exposed to strong light for 15 min. Reverse transcription (RT) followed by quantitative PCR (qPCR) was performed for mRNA extracted from the light-pulsed samples. The RT-qPCR data show that light induction of the *al-3* gene became undetectable in *wc-1^K638A^*, while that of *sub-1* increased prominently in the same mutant (Fig. 4). *al-3* expression was evidently repressed in *wc-1^K694A^*, *wc-2^K244A^*, *wc-2^K417A^*, and *wc-2^K485A^*. Light-activated expression of *ncu06597*, *ncu09635*, *ncu01107*, *ncu02765*, and *ncu00309* (38) was dramatically elevated in *wc-1^K685A^* relative to that in WT (Fig. 4). These data here suggest that acetylation of primary photoreceptors, WC-1 and WC-2, differentially regulates acute light responses in the organism.

**Figure 4.**
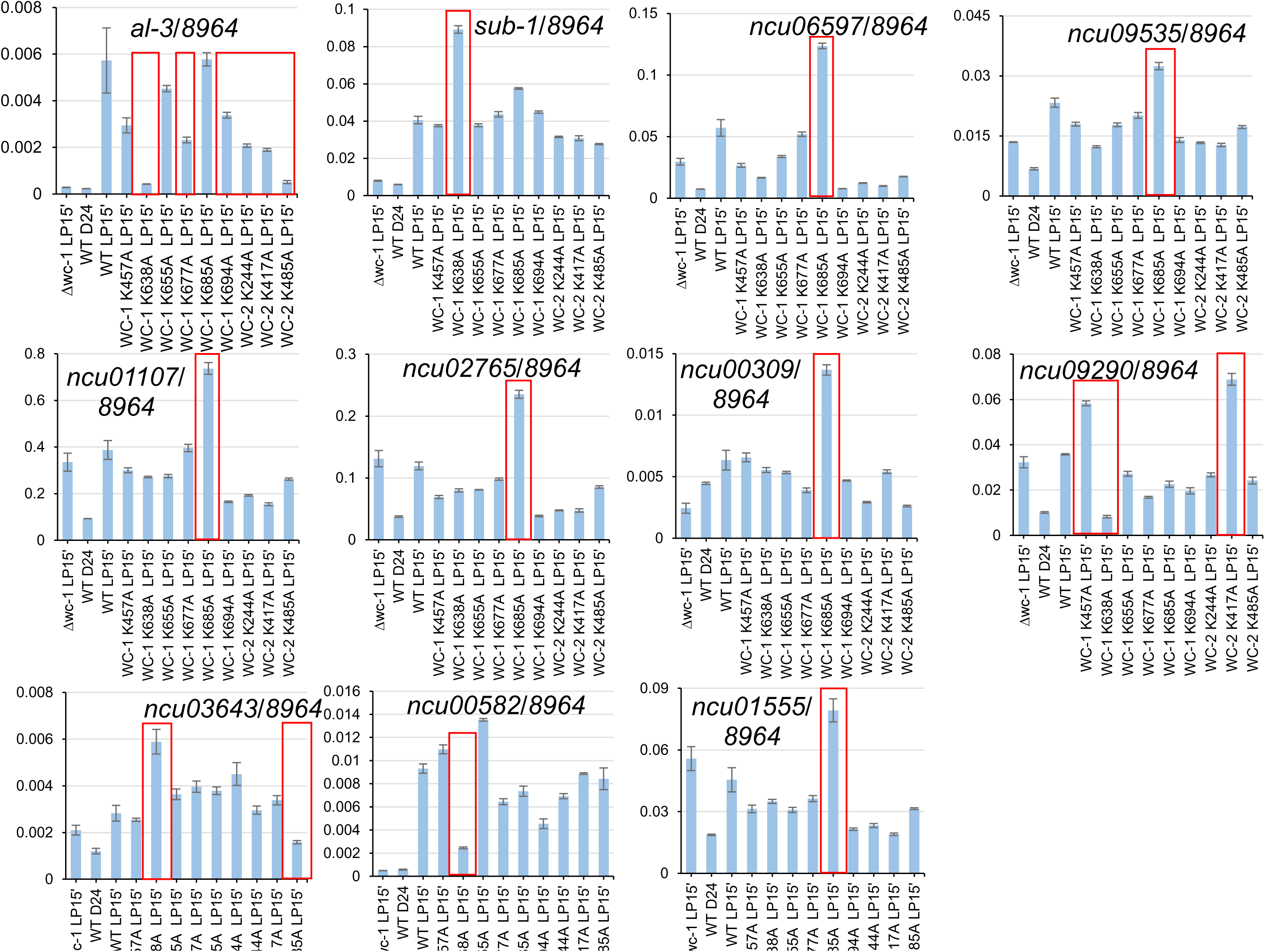
Acute light responses in *wc-1* acetylation mutants. *Δwc-1* and WT serve as negative controls for the light response test. All the strains listed were cultured in the dark for 24 hrs and given a 15-min light pulse except for WT that was only exposed to light. RNA was extracted from the samples, reverse transcription (RT) was performed, and quantitative PCR (qPCR) was conducted with gene-specific primer sets. Red boxes denoted genes with a reproducible change of expression in mutants versus WT. The data presented in the figure come from one experiment, while data obtained from the other independent biological repeat were deposited in the Supporting Fig. 5.

### Identification of mono-methylation sites on WC-1

From the mass-spectrometry data of WC-1 and WC-2, we also identified nine mono-methylation sites on WC-1 (Fig. 5A and Supporting Fig. 3) with the coverage of 51.4% for WC-1 and 46.7% for WC-2 (Supporting Fig. 4). We could not retrieve any di- or tri-methylated, or ubiquitinated peptides from WC-1 and WC-2, which might reflect the low abundance or absence of those modifications on WCC in the cell. The data here clearly indicate that WC-1 is also modified by at least two additional types of PTMs other than phosphorylation.

**Figure 5.**
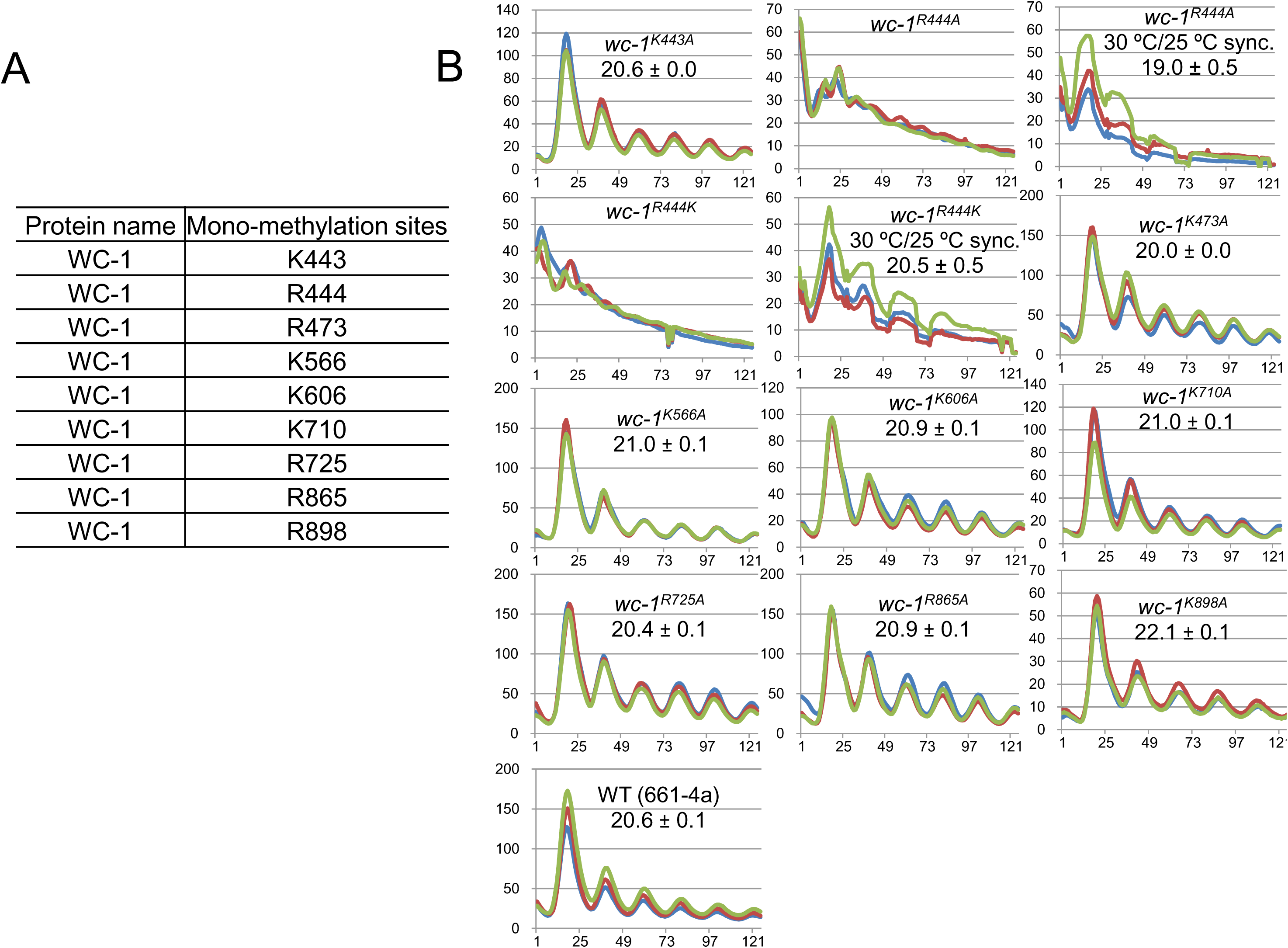
Identification and mutagenesis studies of mono-methylation sites on WC-1 (**A**) Identification of mono-methylation sites on WC-1. (**B**) Luciferase analysis of *wc-1* mutants bearing alanine mutants to the mono-methylation sites as stated. Strains were grown at 25 °C or 30 °C as marked plus light overnight and transferred to the dark at 25 °C. Bioluminescence signals from the dark-grown strains were tracked hourly. Three replicates as shown in three colors are represented with time (in hours) as the x-axis and the signal intensity (arbitrary units) as the y-axis. Period length was calculated from the three replicates and displayed as the average +/- the SEM. All *wc-1* mutants were appended by a V5 tag at the native locus.

### Mutagenesis analysis of mono-methylation sites on WC-1

Like the minor effects of WCC acetylation on the clock (Fig. 1B), most mono-methylation sites (Fig. 5A) when individually mutated to alanine, did not cause evident period alterations, with the exception of R444 (Fig. 5B). Either R444A or R444K caused arrhythmicity, which might be due to the mutation at R444 interfering with the function of the LOV domain (aa 389-506) in light-medicated synchronization of the clock, similar to what was noticed in *wc-1^ΔLOV^* (39). To synchronize the endogenous clock in a way independent of light, *wc-1^R444A^* and *wc-1^R444K^* mutants were grown at 30 °C (still in the light) for ∼ 24 hrs and then moved to the dark at 25°C for recording WCC circadian activity via the *frq C-box*-driven luciferase. The luciferase data reveal that both mutants displayed a robust rhythm with a roughly WT period length (Fig. 5B). The data here are consistent with R444 mutations interfering with light signaling through the LOV domain but not impacting the clock, and suggest that like acetylation, mono-methylation on WCC is also not a determinant of the period length.

## Discussion

In this study, we identified acetylation and mono-methylation sites on WC-1 and WC-2, thereby extending beyond phosphorylation the PTMs found on *Neurospora* clock proteins. Unexpectedly, mutating most of these modified residues individually did not lead to strong period changes, and biochemical analysis did not detect acetylation on WCC, suggesting that acetylation may be not highly penetrant in the WCC population. In the mammalian circadian system, multiple types of PTM events have been documented on the circadian positive element proteins Bmal1/CLOCK, which impact their circadian activity to variable extents (see Introduction). CLOCK has been reported to possess acetyltransferase activity, and one of its substrates is also its interacting partner, Bmal1, whose acetylation at K537 facilitates Cryptochrome (Cry)-mediated repression of the Bmal1/ CLOCK complex (23). However, there are contradictions regarding the timing and the biological relevance of K537 acetylation as well as the acetyltransferase in mediating the modification (see Introduction). In contrast to acetylation on Bmal1/CLOCK, elimination of acetylation on WCC, however, did not result in a pronounced period alteration, suggesting that regulatory acetylation of circadian positive-element proteins is not a conserved mechanism across eukaryotic circadian systems. In the *Drosophila* clock, CYCLE (CYC), the functional ortholog of BMAL1, consists of essentially only the bHLH (basic Helix-Loop-Helix) and PASA/B (Per-Arnt-Sim A/B) domains without an extended C-terminal tail where K537 of Bmal1 is located, so CYC even lacks the acetylation site, K537, of BMAL1, which means that there is little evidence for conservation of regulation involving acetylation even across all animal clocks, much less from animals to fungi. Although multiple PTMs have been reported on Bmal1/CLOCK, which PTM plays a dominant role in dictating the clock operation remains unclear. In *Neurospora*, however, phosphorylation has been shown to be the major mechanism in tuning the circadian activity of WCC (13–15). Especially in a recent study (14), disruption of a small number among over 95 phosphosites that were mapped on WC-1 and WC-2 resulted in constitutively active WCC driving transcription of *frq* and *ccgs* (circadian-controlled genes), indicating that phosphorylation is the principal determinant for WCC repression. WCC serves as the primary photoreceptor for mediating light-induced gene expression genome-wide and also for resetting the core clock through activating *frq* transcription via the *pLRE* element (see Introduction). The RT-qPCR data (Fig. 4) reveal roles of individual acetylation on WCC in regulating acute light responses. Elimination of acetylation at either K638 or K685 causes varying effects on the expression of different light responsive genes, reflecting that acetylation at one individual residue may only modulate expression of a subset of WCC-controlled genes. K638 and K685 are located near PASA and PASB domains of WC-1 (Fig. 6) but are quite distant from the LOV domain, so the impact of the modification may be related to the overall structure of light-activated WCC instead of the formation of the WCC dimer.

**Figure 6.**
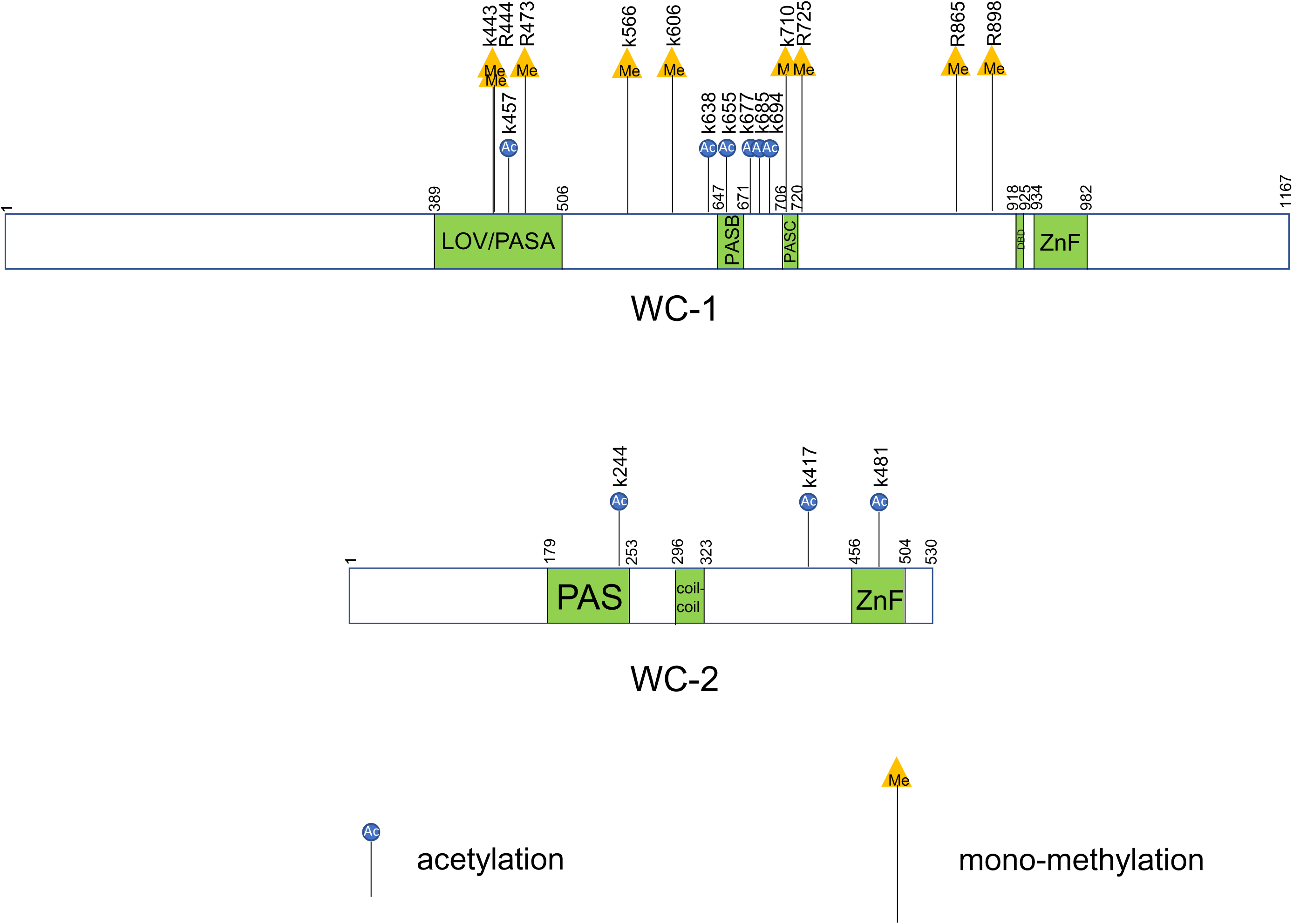
Summary of acetylation and mono-methylation sites identified on WC-1 and WC-2. A schematic diagram of WC-1 and WC-2 shows positions of acetylated and mono-methylated residues identified in this study. WC-1 (1,167 amino acids) contains LOV (light, oxygen, and voltage) (also known as PAS[Per-Arnt-Sim]A), two additional PAS (PASB and PASC), DBD (defective in DNA binding), and ZnF (zinc-finger) DNA-binding domains. WC-2 (530 amino acids) possesses PAS, coiled-coil, and ZnF domains. Blue circles that were labelled with “Ac” inside represent acetylation sites; orange triangles with “Me” inside mean mono-methylation residues. Above the blue circles and orange triangles, vertical letters and their following numbers annotate positions of the modified residues (either a lysine [K] or an arginine [R]).

## Experimental Procedures

### Neurospora strains

*Neurospora* strain 661-4a (*ras-1^bd^*, *A*, *his-3*::*C-box*-driven *luciferase*), WT in luciferase assays, harbors the *C-box* DNA element from the *frq* promoter fused to the codon-optimized firefly *luciferase* gene (transcriptional fusion) at the *his-3* locus(40). *wc-1* and *wc-2* mutants were engineered using a published method relying on yeast homologous recombination-based integration of PCR fragments (33): Restriction-digested shuttle vectors *-1-6* (33) and *-2-1* (14) for *wc-1* and *wc-2* respectively were recombined with PCR (Thermo Fisher Scientific, Catalog # F549S) products amplified with primers bearing point mutations of *wc-1* or *wc-2*. Four primer pairs functioned as flanks in homologous recombination in a *Saccharomyces cerevisiae* strain FY834. All introduced point mutations at target sites were verified by Sanger sequencing in the Dartmouth Core facility with *wc-1*- or *wc-2*-specific primers. All these *wc-1* and *wc-2* mutants were targeted at the native loci. Plasmids bearing mutations were cut with *Ase*I (NEB, Catalog # R0526S) and *Ssp*I (NEB, Catalog # R0132S) and then purified with the QIAquick PCR Purification Kit (Qiagen, Catalog # 28104) for *Neurospora* transformation. *Neurospora* transformation via electroporation (settings: 1,500 V, 600 Ω, and 25 μF) was performed using an electroporator (BTX, Model # ECM630) as previously described (21). The recipient strains for transforming the mutant constructs are Δ*wc-1*::*hph*; Δ*mus-52*::*hph*; *ras-1^bd^* for *wc-1* and Δ*wc-2*::*hph*; Δ*mus-52*::*hph*; *ras-1^bd^* for *wc-2* (14). All *wc-1* and *wc-2* mutants made in this research bear the *ras-1^bd^* mutation and *frq C-box*-driven codon-optimized firefly *luciferase* gene at the *his-3* locus for circadian phenotype analyses, and all mutated WC-1 contain a V5 tag at the C termini for biochemical assays.

### Growth media

For vegetative growth, strains were cultured on complete-medium slants bearing 1 x Vogel’s, 1.6% glycerol, 0.025% casein hydrolysate, 0.5% yeast extract, 0.5% malt extract, and 1.5% agar (41). Sexual crosses between two *Neurospora* strains were conducted at 25 °C in the light on Westergaard’s agar plates containing 1 x Westergaard’s salts, 2% sucrose, 50 ng/mL biotin, and 1.5% agar (42). The Liquid Culture Medium (LCM) contains 1 x Vogel’s, 0.5% arginine, 50 ng/mL biotin, and 2% glucose.

### Protein extraction, immunoprecipitation (IP), and Western blot

Protein extraction, IP, and subsequent WB were completed as previously documented (43, 44). LCM was used in culturing *Neurospora* for IP and WB; cultures were shaking at 125 rpm in the light at 25 °C overnight or as indicated. Vacuum-dried *Neurospora* tissue was frozen immediately in liquid nitrogen and ground to a fine powder with a ceramic mortar & pestle. 10 mL of protein-extraction buffer (50 mM HEPES [pH 7.4], 137 mM NaCl, 10% glycerol, 0.4% NP-40) was mixed with one tablet of cOmplete, Mini, EDTA-free Protease Inhibitor Cocktail (Roche, Catalog # 04693159001).The inhibitors-containing buffer was added to the ground powder followed by vortexing at top speed for 10 sec and then resting on ice for another 10 sec, which was repeated for a total of 2 min prior to incubation on ice for another ten min. Thereafter, centrifugation (12,000-rpm centrifugation at 4 °C for 10 min) was done in order to remove insoluble debris.

For IP with *Neurospora* lysate, two mgs of extracts (precleaned by centrifugation at 12,000-rpm and 4 °C for 10 min) were incubated with 20 μL of FLAG resin (Sigma-Aldrich, Catalog # A2220) followed by rotating at 4 °C for 2 hrs. The protein-bound resin was washed (inverted ten times) twice each with one mL of the protein-extraction buffer bearing protease inhibitors and afterwards eluted with 100 μL of 5 × SDS sample buffer being heated at 99 °C for 5 min. Ten out of the 100-μL IP product were loaded per lane in a gel.

For WB, 15 μgs of centrifugation (12,000 rpm at 4 °C for 10 min)-cleared whole cell lysate were loaded per lane in a commercial 3-8% 1.5-mm x 15- well Tris-Acetate SDS gel (Thermo Fisher Scientific, Catalog # EA03785BOX), electrophoresing with 1 x NuPAGE Tris-Acetate SDS Running Buffer (Thermo Fisher Scientific, Catalog # LA0041). Applications of custom rabbit FRQ, FRH, WC-1, and WC-2 antibodies in WB have been reported previously (39). Antibodies against acetylated lysines include Acetylated-Lysine Mouse mAb (Ac-K-103) (Cell Signaling Technology [CST], Catalog # 9681), Acetylated Lysine Monoclonal Antibody (1C6) (Invitrogen, Catalog # MA1-2021), and Ac-lysine Antibody (AKL5C1) (Santa Cruz Biotechnology, Catalog # sc-32268), each of which was employed in Western blotting at a dilution of 1:1,000.

### Identification of acetylation and mono-methylation sites on WC-1 and WC-2 by mass spectrometry

Purified WC-1 and WC-2 were processed and analyzed by LC-MS/MS (liquid chromatography-coupled tandem mass spectrometry) as previously described (14, 33). Raw data were searched against a custom database containing WC-1 and WC-2 sequences with a peptide mass tolerance of 1 Da, static modification carbamidomethyl (C) +57.02146, and dynamic modifications oxidation (M) +15.99491 and acetylation (K) +42.010565 or monomethylation (K, R) +14.01565.

### Luciferase-reporter assays

Luciferase assays were done following former methods (45, 46). 96-black well plates (0.8 mL of the luciferase-assay medium in each well) were inoculated with conidial suspension (conidia in water), and the inoculants were grown at 25 °C in the presence of light for 16–24 hrs and then transferred to the dark at 25 °C for monitoring light production. Medium used for luciferase assays bore 1 x Vogel’s salts, 0.17% arginine, 1.5% Bacto Agar, 50 ng/mL biotin, 0.1% glucose, and 12.5 μM luciferin (GoldBio, Catalog # LUCK-2G). Bioluminescence signals were tracked with a CCD (charged-coupled device) camera every hour, raw data were acquired with ImageJ and a custom macro, and period lengths were manually determined. Raw data from three or four replicates were shown in the Figures, and time (in hours) on the x-axis was plotted against the signal intensity (arbitrary units) on the y-axis. WT used in luciferase assays was 661-4a (*ras-1^bd^*, *A*) with the codon-optimized firefly *luciferase* gene driven by the *frq C-box* at the *his-3* locus.

### RT-quantitative PCR

TRIzol reagent (Life Technologies) and SuperScript III first strand synthesis kit (Life Technologies) were used to isolate *Neurospora* mRNA and to synthesize cDNA (from 3 µg of mRNA) respectively (38). Real-time PCR was conducted with iTaq Universal SYBR green Supermix (BIO-RAD, Catalog # 1725121) using an ABI 7500 Fast system. Sequences of primers against *wcc* target genes were obtained from prior publications (38, 45, 46). *ncu08964* served as an internal normalizer for RT-qPCR, which is constantly expressed under different nutrient or circadian conditions.

## Acknowledgements

This research was supported by National Institutes-of-Health (NIH) grants awarded to Jay C. Dunlap (R35GM118021), Arminja N. Kettenbach (R35GM119455), and Scott A. Gerber (R35GM145596).

## Data availability

Mutants made in this study are available upon request. All data that were used to draw conclusions in the article have been provided in the figures.

## Author Contributions

Conceptualization, B.W. and J.C.D.; methodology, B.W. M.E.E. X.Y.Z. S.A.G. and A.N.K.; software, B.W. M.E.E. and A.N.K.; formal analysis, B.W. M.E.E. and A.N.K.; investigation, B.W. M.E.E. X.Y.Z. S.A.G. and A.N.K.; resources, J.C.D. A.N.K. and S.A.G.; writing—original draft preparation, B.W. and A.N.K.; writing—review and editing, J.C.D.; funding acquisition, J.C.D. A.N.K. and S.A.G.

## Figure legends

**Supporting Table S1** List of identified acetylated and mono-methylated peptides derived from WC-1 and WC-2 by mass spectrometry. In the table, “*”, “∼”, and “!” indicates oxidation, acetylation, and mono-methylation, respectively.

**Supporting Fig. 1** Mass spectrometry spectra of acetylated peptides from WC-1 and WC-2. WC-1 and WC-2 were purified from cultures that grew in the light at 25 °C. Digestion of WC-1 and WC-2 was carried out with either proteinase K or trypsin as indicated in the Supporting Table 1. The spectra of the acetylated peptides were generated using IPSA (Interactive Peptide Spectra Annotator) (47). The x-axes in the charts are the ratio of mass to charge (m / z), while the y-axes are relative signal intensities (labelled as “relative abundance [%]”). Red arrowheads point to residues that are acetylated. In the annotated spectra, lower case “c” means carbamidomethylcysteine; lower case “m” represents oxidated methionine; lower case “k” indicates acetylated lysine. Nine acetylated peptide species of WC-1 and WC-2 were discovered in the analysis, and all the acetylated peptides of WC-1 and WC-2 have been deposited in the Supporting Table 1.

**Supporting Fig. 2** Coverage maps of WC-1 and WC-2 in acetylation analysis. Residues covered by mass spectrometry are in red, while undetected residues are in grey.

**Supporting Fig. 3** Mass spectrometry spectra of mono-methylated and unmethylated peptides from WC-1. WC-1 and WC-2 were isolated from light-grown cultures (at 25 °C), and digested proteinase K or trypsin as listed in the Supporting Table 1. The spectra of the mono-methylated peptides were produced by IPSA (Interactive Peptide Spectra Annotator) (47). In the spectra, lower case “c” denotes carbamidomethylcysteine; lower case “m” means oxidated methionine; lower case “r” or “k” represents mono-methylated arginine or lysine. The x-axes in the charts represent the mass to charge ratio (m / z), while the y-axes mean relative abundance [%]. Ten mono-methylated peptide species of WC-1 were found in the analysis, and all the mono-methylated WC-1 peptides have been put in the Supporting Table 1. Red arrowheads indicate WC-1 residues that were found to be modified by mono-methylation. Two types of mono-methylated peptides have been identified at the residue K443 of WC-1.

**Supporting Fig. 4** Coverage maps of WC-1 and WC-2 in mono-methylation analysis. Mass spectrometry-identified residues were labeled in red and the rest are in grey.

**Supporting Fig. 5** RT-qPCR data obtained from another independent biological repeat (relative to Fig. 4) using *wc-1* and *wc-2* mutants (Fig. 1B) bearing alanine mutations to individual acetylation sites as indicated.

## References

1. Bell-Pedersen, D., Cassone, V. M., Earnest, D. J., Golden, S. S., Hardin, P. E., Thomas, T. L., and Zoran, M. J. (2005) Circadian rhythms from multiple oscillators: lessons from diverse organisms. Nat Rev Genet. 6, 544–556

2. Roenneberg, T., and Merrow, M. (2016) The Circadian Clock and Human Health. Current Biology. 26, R432–R443

3. Shen, L., Su, Z., Yang, K., Wu, C., Becker, T., Bell-Pedersen, D., Zhang, J., and Sachs, M. S. (2021) Structure of the translating *Neurospora* ribosome arrested by cycloheximide. Proc. Natl. Acad. Sci. U.S.A. 118, e2111862118

4. Castillo, K. D., Wu, C., Ding, Z., Lopez-Garcia, O. K., Rowlinson, E., Sachs, M. S., and Bell-Pedersen, D. (2022) A circadian clock translational control mechanism targets specific mRNAs to cytoplasmic messenger ribonucleoprotein granules. Cell Reports. 41, 111879

5. Zhang, C., Tian, Y., Song, S., Zhang, L., Dang, Y., and He, Q. (2022) H3K56 deacetylation and H2A.Z deposition are required for aberrant heterochromatin spreading. Nucleic Acids Research. 50, 3852–3866

6. Froehlich, A. C., Liu, Y., Loros, J. J., and Dunlap, J. C. (2002) White Collar-1, a Circadian Blue Light Photoreceptor, Binding to the *frequency* Promoter. Science. 297, 815–819

7. Froehlich, A. C., Loros, J. J., and Dunlap, J. C. (2003) Rhythmic binding of a WHITE COLLAR-containing complex to the *frequency* promoter is inhibited by FREQUENCY. Proc. Natl. Acad. Sci. U.S.A. 100, 5914–5919

8. Froehlich, A. C., Liu, Y., Loros, J. J., and Dunlap, J. C. (2002) White Collar-1, a Circadian Blue Light Photoreceptor, Binding to the *frequency* Promoter. Science. 297, 815–819

9. He, Q., Cheng, P., Yang, Y., Wang, L., Gardner, K. H., and Liu, Y. (2002) White Collar-1, a DNA Binding Transcription Factor and a Light Sensor. Science. 297, 840–843

10. Shi, M., Collett, M., Loros, J. J., and Dunlap, J. C. (2010) FRQ-Interacting RNA Helicase Mediates Negative and Positive Feedback in the Neurospora Circadian Clock. Genetics. 184, 351–361

11. Cheng, P., He, Q., He, Q., Wang, L., and Liu, Y. (2005) Regulation of the *Neurospora* circadian clock by an RNA helicase. Genes Dev. 19, 234–241

12. Jankowski, M. S., Griffith, D., Shastry, D. G., Pelham, J. F., Ginell, G. M., Thomas, J., Karande, P., Holehouse, A. S., and Hurley, J. M. (2022) The formation of a fuzzy complex in the negative arm regulates the robustness of the circadian clock, bioRxiv, 10.1101/2022.01.04.474980

13. He, Q., Cha, J., He, Q., Lee, H.-C., Yang, Y., and Liu, Y. (2006) CKI and CKII mediate the FREQUENCY-dependent phosphorylation of the WHITE COLLAR complex to close the *Neurospora* circadian negative feedback loop. Genes Dev. 20, 2552–2565

14. Wang, B., Kettenbach, A. N., Zhou, X., Loros, J. J., and Dunlap, J. C. (2019) The Phospho-Code Determining Circadian Feedback Loop Closure and Output in Neurospora. Molecular Cell. 74, 771–784.e3

15. Schafmeier, T., Haase, A., Káldi, K., Scholz, J., Fuchs, M., and Brunner, M. (2005) Transcriptional Feedback of Neurospora Circadian Clock Gene by Phosphorylation-Dependent Inactivation of Its Transcription Factor. Cell. 122, 235–246

16. Sancar, G., Sancar, C., Brunner, M., and Schafmeier, T. (2009) Activity of the circadian transcription factor White Collar Complex is modulated by phosphorylation of SP-motifs. FEBS Letters. 583, 1833–1840

17. Garceau, N. Y., Liu, Y., Loros, J. J., and Dunlap, J. C. (1997) Alternative Initiation of Translation and Time-Specific Phosphorylation Yield Multiple Forms of the Essential Clock Protein FREQUENCY. Cell. 89, 469–476

18. Yang, Y., Cheng, P., and Liu, Y. (2002) Regulation of the *Neurospora* circadian clock by casein kinase II. Genes Dev. 16, 994–1006

19. Baker, C. L., Kettenbach, A. N., Loros, J. J., Gerber, S. A., and Dunlap, J. C. (2009) Quantitative Proteomics Reveals a Dynamic Interactome and Phase-Specific Phosphorylation in the Neurospora Circadian Clock. Molecular Cell. 34, 354–363

20. Tang, C.-T., Li, S., Long, C., Cha, J., Huang, G., Li, L., Chen, S., and Liu, Y. (2009) Setting the pace of the *Neurospora* circadian clock by multiple independent FRQ phosphorylation events. Proc. Natl. Acad. Sci. U.S.A. 106, 10722–10727

21. Wang, B., Stevenson, E.-L., and Dunlap, J. C. (2023) Functional analysis of 110 phosphorylation sites on the circadian clock protein FRQ identifies clusters determining period length and temperature compensation. G3 Genes|Genomes|Genetics. 13, jkac334

22. Partch, C. L., Green, C. B., and Takahashi, J. S. (2014) Molecular architecture of the mammalian circadian clock. Trends in Cell Biology. 24, 90–99

23. Hirayama, J., Sahar, S., Grimaldi, B., Tamaru, T., Takamatsu, K., Nakahata, Y., and Sassone-Corsi, P. (2007) CLOCK-mediated acetylation of BMAL1 controls circadian function. Nature. 450, 1086– 1090

24. Petkau, N., Budak, H., Zhou, X., Oster, H., and Eichele, G. (2019) Acetylation of BMAL1 by TIP60 controls BRD4-P-TEFb recruitment to circadian promoters. eLife. 8, e43235

25. Cardone, L., Hirayama, J., Giordano, F., Tamaru, T., Palvimo, J. J., and Sassone-Corsi, P. (2005) Circadian Clock Control by SUMOylation of BMAL1. Science. 309, 1390–1394

26. Lee, J., Lee, Y., Lee, M. J., Park, E., Kang, S. H., Chung, C. H., Lee, K. H., and Kim, K. (2008) Dual Modification of BMAL1 by SUMO2/3 and Ubiquitin Promotes Circadian Activation of the CLOCK/BMAL1 Complex. Mol Cell Biol. 28, 6056–6065

27. Stojkovic, K., Wing, S. S., and Cermakian, N. (2014) A central role for ubiquitination within a circadian clock protein modification code. Front. Mol. Neurosci. 10.3389/fnmol.2014.00069

28. Jiang, H., Wang, X., Ma, J., and Xu, G. (2023) The fine-tuned crosstalk between lysine acetylation and the circadian rhythm. Biochimica et Biophysica Acta (BBA) - Gene Regulatory Mechanisms. 1866, 194958

29. Nakahata, Y., Kaluzova, M., Grimaldi, B., Sahar, S., Hirayama, J., Chen, D., Guarente, L. P., and Sassone-Corsi, P. (2008) The NAD+- Dependent Deacetylase SIRT1 Modulates CLOCK-Mediated Chromatin Remodeling and Circadian Control. Cell. 134, 329–340

30. Nakahata, Y., Sahar, S., Astarita, G., Kaluzova, M., and Sassone-Corsi, P. (2009) Circadian Control of the NAD ^+^ Salvage Pathway by CLOCK-SIRT1. Science. 324, 654–657

31. Brenna, A., Grimaldi, B., Filetici, P., and Ballario, P. (2012) Physical association of the WC-1 photoreceptor and the histone acetyltransferase NGF-1 is required for blue light signal transduction in *Neurospora crassa*. MBoC. 23, 3863–3872

32. Collett, M. A., Dunlap, J. C., and Loros, J. J. (2001) Circadian Clock-Specific Roles for the Light Response Protein WHITE COLLAR-2. Molecular and Cellular Biology. 21, 2619–2628

33. Wang, B., Kettenbach, A. N., Gerber, S. A., Loros, J. J., and Dunlap, J. C. (2014) Neurospora WC-1 Recruits SWI/SNF to Remodel frequency and Initiate a Circadian Cycle. PLoS Genet. 10, e1004599

34. Cheng, P., Yang, Y., Gardner, K. H., and Liu, Y. (2002) PAS Domain-Mediated WC-1/WC-2 Interaction Is Essential for Maintaining the Steady-State Level of WC-1 and the Function of Both Proteins in Circadian Clock and Light Responses of *Neurospora*. Mol Cell Biol. 22, 517–524

35. Xiong, L., Adhvaryu, K. K., Selker, E. U., and Wang, Y. (2010) Mapping of lysine methylation and acetylation in core histones of Neurospora crassa. Biochemistry. 49, 5236–5243

36. Adhvaryu, K. K., Berge, E., Tamaru, H., Freitag, M., and Selker, E. U. (2011) Substitutions in the Amino-Terminal Tail of Neurospora Histone H3 Have Varied Effects on DNA Methylation. PLoS Genet. 7, e1002423

37. Cheng, P., Yang, Y., and Liu, Y. (2001) Interlocked feedback loops contribute to the robustness of the *Neurospora* circadian clock. Proc. Natl. Acad. Sci. U.S.A. 98, 7408–7413

38. Chen, C.-H., Ringelberg, C. S., Gross, R. H., Dunlap, J. C., and Loros, J. J. (2009) Genome-wide analysis of light-inducible responses reveals hierarchical light signalling in Neurospora. EMBO J. 28, 1029–1042

39. Wang, B., Zhou, X., Loros, J. J., and Dunlap, J. C. (2016) Alternative Use of DNA Binding Domains by the *Neurospora* White Collar Complex Dictates Circadian Regulation and Light Responses. Molecular and Cellular Biology. 36, 781–793

40. Larrondo, L. F., Olivares-Yañez, C., Baker, C. L., Loros, J. J., and Dunlap, J. C. (2015) Decoupling circadian clock protein turnover from circadian period determination. Science. 347, 1257277

41. Vogel, H.J. (1956) A Convenient Growth Medium for Neurospora crassa, Microbial Genetics Bulletin 13, 42–47

42. Westergaard, M., and Mitchell, H. K. (1947) NEUROSPORA V. A SYNTHETIC MEDIUM FAVORING SEXUAL REPRODUCTION. American Journal of Botany. 34, 573–577

43. Wang, B., and Dunlap, J. C. (2023) Domains required for the interaction of the central negative element FRQ with its transcriptional activator WCC within the core circadian clock of Neurospora. Journal of Biological Chemistry. 299, 104850

44. Wang, B., Zhou, X., Kettenbach, A. N., Mitchell, H. D., Markillie, L. M., Loros, J. J., and Dunlap, J. C. (2023) A crucial role for dynamic expression of components encoding the negative arm of the circadian clock. Nat Commun. 14, 3371

45. Zhou, X., Wang, B., Emerson, J. M., Ringelberg, C. S., Gerber, S. A., Loros, J. J., and Dunlap, J. C. (2018) A HAD family phosphatase CSP-6 regulates the circadian output pathway in Neurospora crassa. PLoS Genet. 14, e1007192

46. Wang, B., Zhou, X., Gerber, S. A., Loros, J. J., and Dunlap, J. C. (2021) Cellular Calcium Levels Influenced by NCA-2 Impact Circadian Period Determination in *Neurospora*. mBio. 12, e01493–21

47. Brademan, D. R., Riley, N. M., Kwiecien, N. W., and Coon, J. J. (2019) Interactive Peptide Spectral Annotator: A Versatile Web-based Tool for Proteomic Applications. Mol Cell Proteomics. 18, S193–S201

