## Supplementary material for "Acetylation of WCC is dispensable for the core circadian clock but differentially regulates acute light responses in *Neurospora*": suppl. figs

### Supporting Figure 1

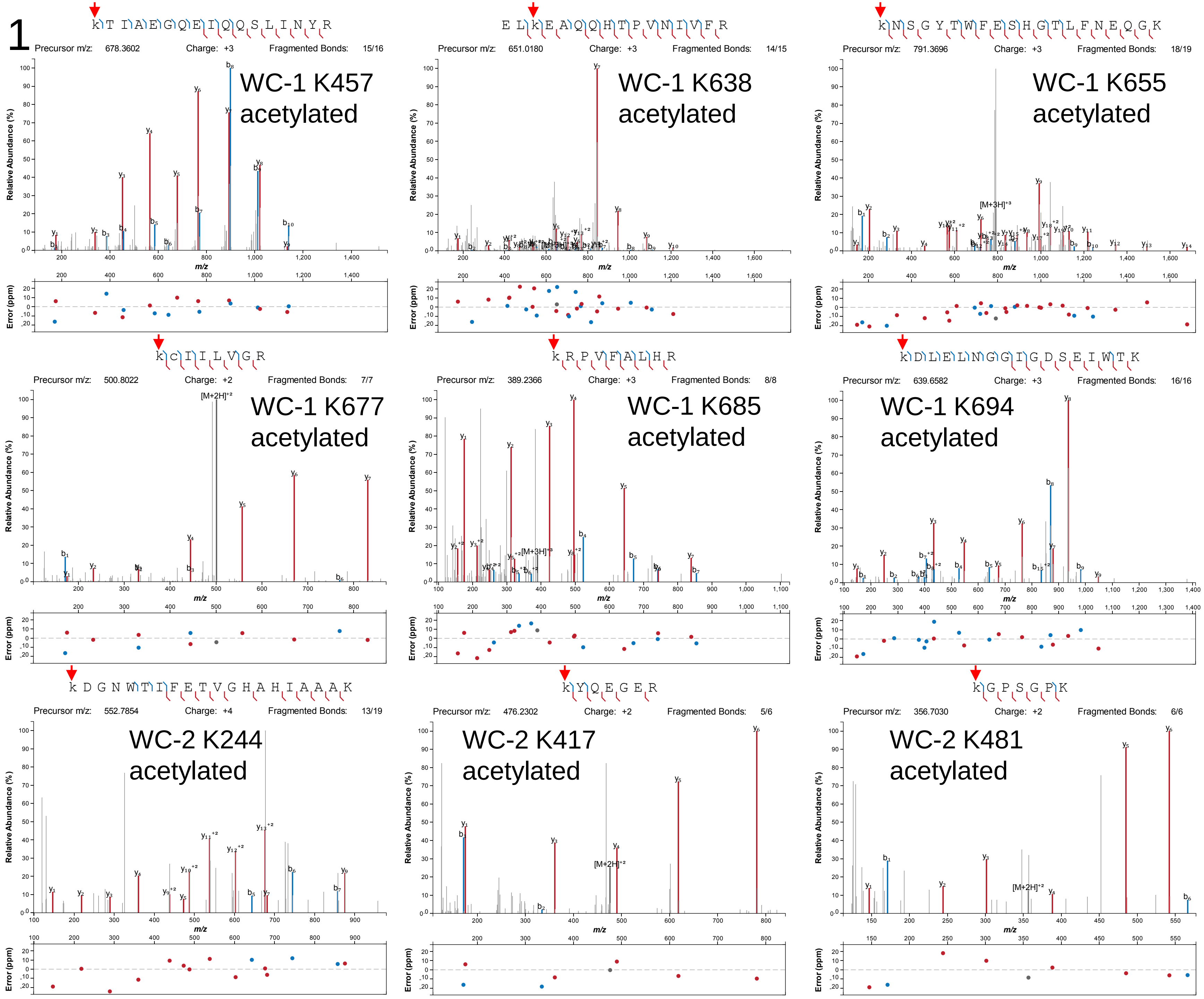

### Supporting Figure 2

#### Mass spectrometry coverage maps for acetylation identification on WC-1 and WC-2

White collar 1 protein OS=Neurospora crassa (strain ATCC 24698 / 74-OR23-1A / CBS 708.71 / DSM 1257 / FGSC 987) GN=wc-1 PE=1 SV=2
Uniprot | UCSC Proteome Browser
Molecular Weight: 127,355

```
MNNNYYGSPL SPEELQHQMH QHQQQQQQQQ QQQQQQQQQQ QQQQQQQQQQ HQHQQQQKTN QHRNAGMMNT PPTTNQGNST IHASDVTMSG GSDSLDEIIQ
QNLDEMHRRR SVPQPYGGQT RRLSMFDYAN PNDGFSDYQL DNMSGNYGDM TGGMGMSGHS SPYAGQNIMA MSDHSGGYSH MSPNVMGNMM TYPNLNMYHS
PPIENPYSSA GLDTIRTDFS MDMNMDSGSV SAASVHPTPG LNKQDDEMMT MEQGFGGGDD ANASHQAQQN MGGLTPAMTP AMTPAMTPGV SNFAQGMATP
VSQDAASTPA TTFQSPSLSA TTQTIRIGPP PPPSVTNAPT PAPFTSTPSG GGASQTKSIY SKSGFDMLRA LWYVASRKDP KLKLGAVDMS CAFVVCDVTL
NDCPIIYVSD NFQNLTGYSR HEIVGRNCRF LQAPDGNVEA GTKREFVENN AVYTLKKTIA EGQEIQQSLI NYRKGGKPFL NLLTMIPIPW DTEEIRYFIG
FQIDLVECPD AIIGQEGNGP MQVNYTHSDI GQYIWTPPTQ KQLEPADGQT LGVDDVSTLL QQCNSKGVAS DWHKQSWDKM LLENADDVVH VLSLKGLFLY
LSPACKKVLE YDASDLVGTS LSSICHPSDI VPVTRELKEA QQHTPVNIVF RIRRKNSGYT WFESHGTLFN EQGKGRKCII LVGRKRPVFA LHRKDLELNG
GIGDSEIWTK VSTSGMFLFV SSNVRSLLDL LPENLQGTSM QDLMRKESRA EFGRTIEKAR KGKIASCKHE VQNKRGQVLQ AYTTFYPGDG GEGQRPTFLL
AQTKLLKASS RTLAPATVTV KNMSPGGVPL SPMKGIQTDS DSNTLMGGMS KSGSSDSTGA MVSARSSAGP GQDAALDADN IFDELKTTRC TSWQYELRQM
EKVNRMLAEE LAQLLSNKKK RKRRKGGGNM VRDCANCHTR NTPEWRRGPS GNRDLCNSCG LRWAKQTGRV SPRTSSRGGN GDSMSKKSNS PSHSSPLHRE
VGNDSPSTTT ATKNSPSLRG SSTTAPGTIT TDSGPAVASS ASGTGSTTIA TSANSAASTV NALGPPATGP SGGSPAQHLP PHLQGTHLNA QAMQRVHQHK
QHQQHQQQHQ QQHQQQHQQQ HQQLQQHQFN PPQSQPLLEG GSGFRGSGME MTSIREEMGE HQQGLSV
```

White collar 2 protein OS=Neurospora crassa (strain ATCC 24698 / 74-OR23-1A / CBS 708.71 / DSM 1257 / FGSC 987) GN=wc-2 PE=1 SV=1
Uniprot | UCSC Proteome Browser
Molecular Weight: 56,840

```
MSHGQPPPGS SMYGFGAMGM GSGMGSGMGS GMGTGMGTGM GTGMSASQMT SDPQDMMSLL DTSVFPGFDG MSMSLDVGDS MSNPFTPVSV PPPLPAGNAG
PSHVGVCGGH GAPDQLFSPD DLIATSMSSA GPMIATPTTT TSGPSGGPSS GGGSTLTEFT KRRNWPAKVV EELQDWEHIL DANGRIKHVS PSVEPLTGYK
PPEIIDLFLR DLIHPDDVGV FTAELNEAIA TGSQLRLFYR FRKKDGNWTI FETVGHAHIA AAKFAPNPQN QSPFCQAVFM MARPYPTKNA GLLDSFLEHK
IENERLKRRI AELRREEQEE QEESHRTWRM SQEGRSDVTP SDDTATQMGM TPFYIPMNAQ ADVMMPPPSQ PASSLNIALT RENLEGIAGS RPDSIREKML
RYEGNHADTI EMLTGLKYQE GERSHGITTG NASPTLIKGD AGIAIPLDRD PRTGEKKKKI KVAEEYVCTD CGTLDSPEWR KGPSGPKTLC NACGLRWAKK
EKKKNANNNN NGGGIGGHND IHTPMGDHMG
```

### Supporting Figure 3

**Annotated spectra**
Lower case c – carbamidomethylcysteine
Lower case m – oxidated methionine
Lower case r or k – monomethylated arginine or lysine, or acetylated lysine

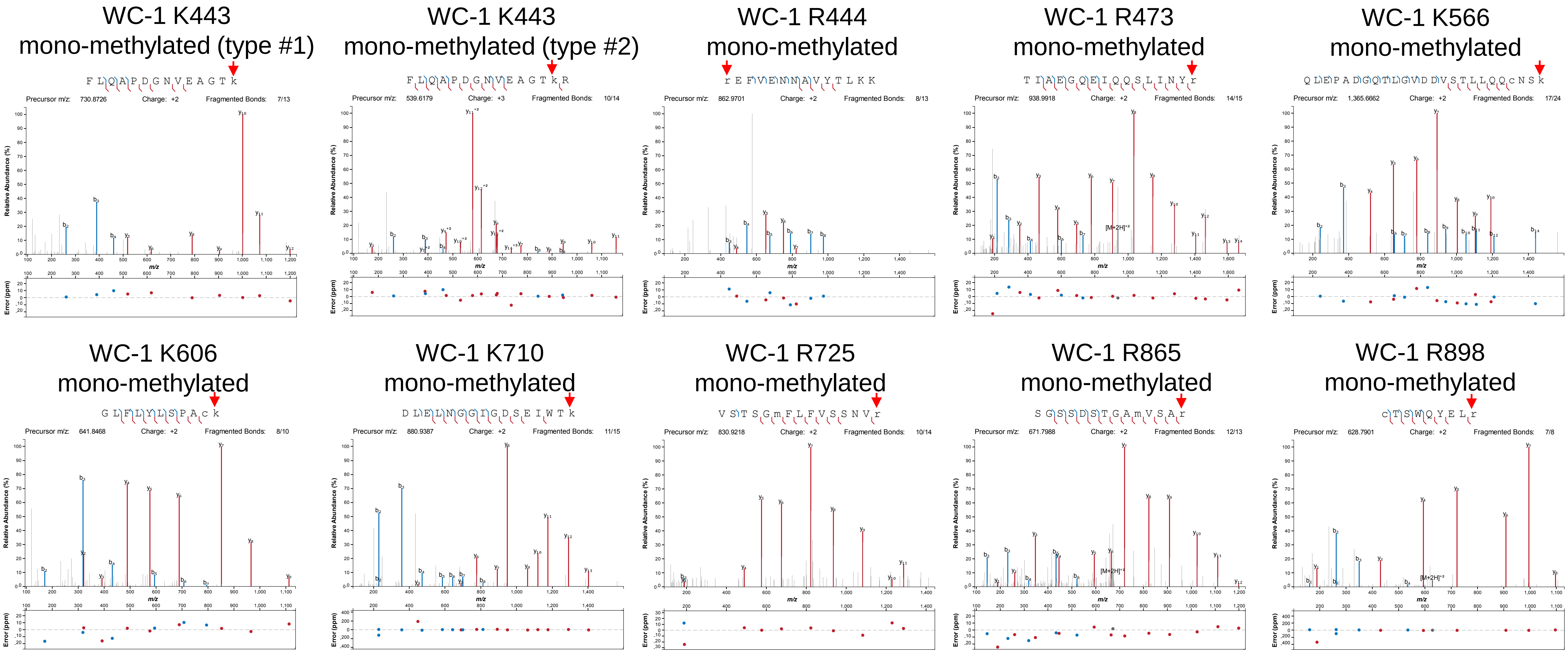

### Supporting Figure 3 continued

**Annotated spectra**
Lower case c – carbamidomethylcysteine
Lower case m – oxidated methionine
Lower case r or k – monomethylated arginine or lysine, or acetylated lysine

K443 non-methyl

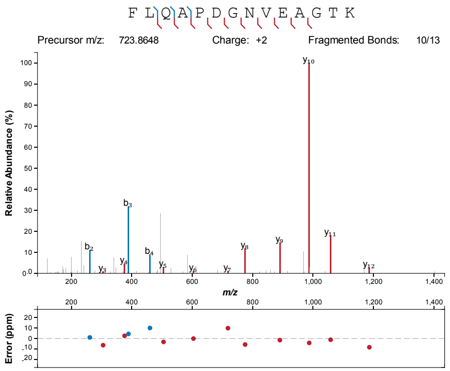


K443 non-methyl

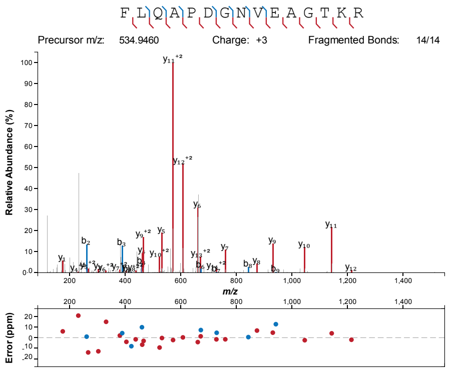


R444 non-methyl

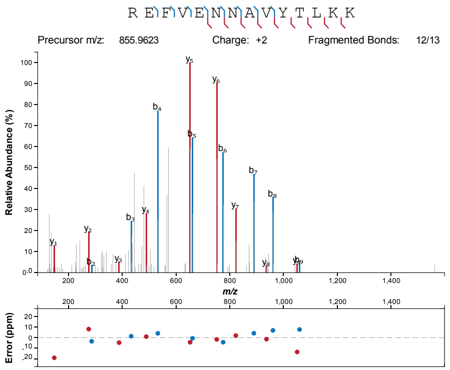


R473 non-methyl

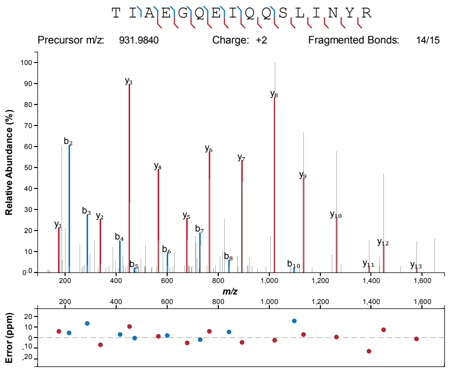


K566 non-methyl

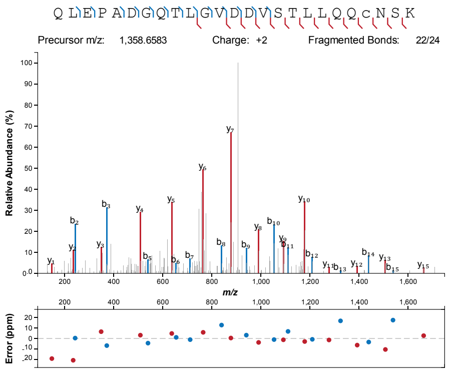


K606 non-methyl

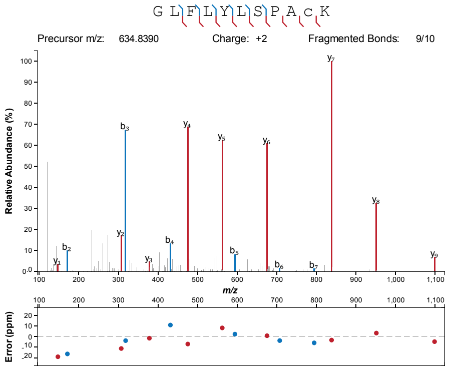


K710 non-methyl

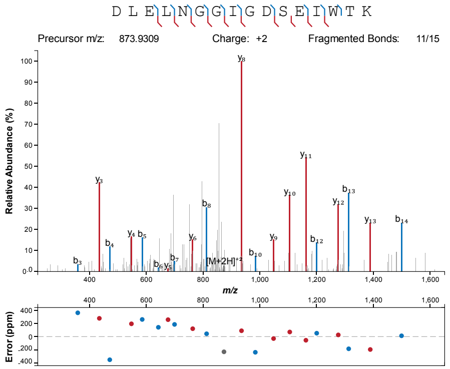


R725 non-methyl

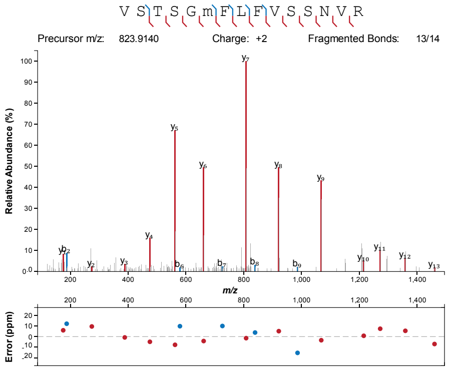


R865 non-methyl

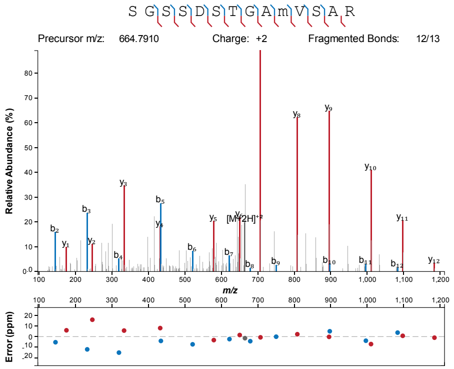


R898 non-methyl

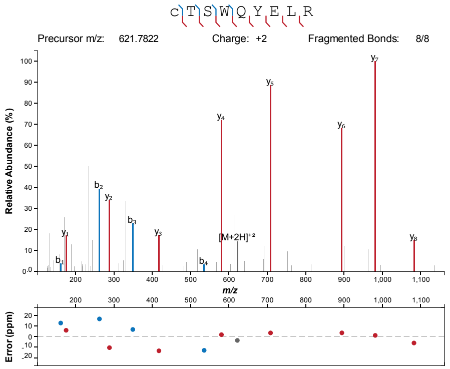

### Supporting Figure 4

#### Mass spectrometry coverage maps for mono-methylation identification on WC-1 and WC-2

White collar 1 protein OS=Neurospora crassa (strain ATCC 24698 / 74-OR23-1A / CBS 708.71 / DSM 1257 / FGSC 987) GN=wc-1 PE=1 SV=2
Uniprot | UCSC Proteome Browser
Molecular Weight: 127,355

```
MNNNYYGSPL SPEELQHQMH QHQQQQQQQQ QQQQQQQQQQ QQQQQQQQQQ HQHQQQQKTN QHRNAGMMNT PPTTNQGNST IHASDVTMSG GSDSLDEIIQ
QNLDEMHRRR SVPQPYGGQT RRLSMFDYAN PNDGFSDYQL DNMSGNYGDM TGGMGMSGHS SPYAGQNIMA MSDHSGGYSH MSPNVMGNMM TYPNLNMYHS
PPIENPYSSA GLDTIRTDFS MDMNMDSGSV SAASVHPTPG LNKQDDEMMT MEQGFGGGDD ANASHQAQQN MGGLTPAMTP AMTPAMTPGV SNFAQGMATP
VSQDAASTPA TTFQSPSLSA TTQTIRIGPP PPPSVTNAPT PAPFTSTPSG GGASQTKSIY SKSGFDMLRA LWYVASRKDP KLKLGAVDMS CAFVVCDVTL
NDCPIIYVSD NFQNLTGYSR HEIVGRNCRF LQAPDGNVEA GTKREFVENN AVYTLKKTIA EGQEIQQSLI NYRKGGKPFL NLLTMIPIPW DTEEIRYFIG
FQIDLVECPD AIIGQEGNGP MQVNYTHSDI GQYIWTPPTQ KQLEPADGQT LGVDDVSTLL QQCNSKGVAS DWHKQSWDKM LLENADDVVH VLSLKGLFLY
LSPACKKVLE YDASDLVGTS LSSICHPSDI VPVTRELKEA QQHTPVNIVF RIRRKNSGYT WFESHGTLFN EQGKGRKCII LVGRKRPVFA LHRKDLELNG
GIGDSEIWTK VSTSGMFLFV SSNVRSLLDL LPENLQGTSM QDLMRKESRA EFGRTIEKAR KGKIASCKHE VQNKRGQVLQ AYTTFYPGDG GEGQRPTFLL
AQTKLLKASS RTLAPATVTV KNMSPGGVPL SPMKGIQTDS DSNTLMGGMS KSGSSDSTGA MVSARSSAGP GQDAALDADN IFDELKTTRC TSWQYELRQM
EKVNRMLAEE LAQLLSNKKK RKRRKGGGNM VRDCANCHTR NTPEWRRGPS GNRDLCNSCG LRWAKQTGRV SPRTSSRGGN GDSMSKKSNS PSHSSPLHRE
VGNDSPSTTT ATKNSPSLRG SSTTAPGTIT TDSGPAVASS ASGTGSTTIA TSANSAASTV NALGPPATGP SGGSPAQHLP PHLQGTHLNA QAMQRVHQHK
QHQQHQQQHQ QQHQQQHQQQ HQQLQQHQFN PPQSQPLLEG GSGFRGSGME MTSIREEMGE HQQGLSV
```

White collar 2 protein OS=Neurospora crassa (strain ATCC 24698 / 74-OR23-1A / CBS 708.71 / DSM 1257 / FGSC 987) GN=wc-2 PE=1 SV=1
Uniprot | UCSC Proteome Browser
Molecular Weight: 56,840

```
MSHGQPPPGS SMYGFGAMGM GSGMGSGMGS GMGTGMGTGM GTGMSASQMT SDPQDMMSLL DTSVFPGFDG MSMSLDVGDS MSNPFTPVSV PPPLPAGNAG
PSHVGVCGGH GAPDQLFSPD DLIATSMSSA GPMIATPTTT TSGPSGGPSS GGGSTLTEFT KRRNWPAKVV EELQDWEHIL DANGRIKHVS PSVEPLTGYK
PPEIIDLFLR DLIHPDDVGV FTAELNEAIA TGSQLRLFYR FRKKDGNWTI FETVGHAHIA AAKFAPNPQN QSPFCQAVFM MARPYPTKNA GLLDSFLEHK
IENERLKRRI AELRREEQEE QEESHRTWRM SQEGRSDVTP SDDTATQMGM TPFYIPMNAQ ADVMMPPPSQ PASSLNIALT RENLEGIAGS RPDSIREKML
RYEGNHADTI EMLTGLKYQE GERSHGITTG NASPTLIKGD AGIAIPLDRD PRTGEKKKKI KVAEEYVCTD CGTLDSPEWR KGPSGPKTLC NACGLRWAKK
EKKKNANNNN NGGGIGGHND IHTPMGDHMG
```

### Supporting Figure 5

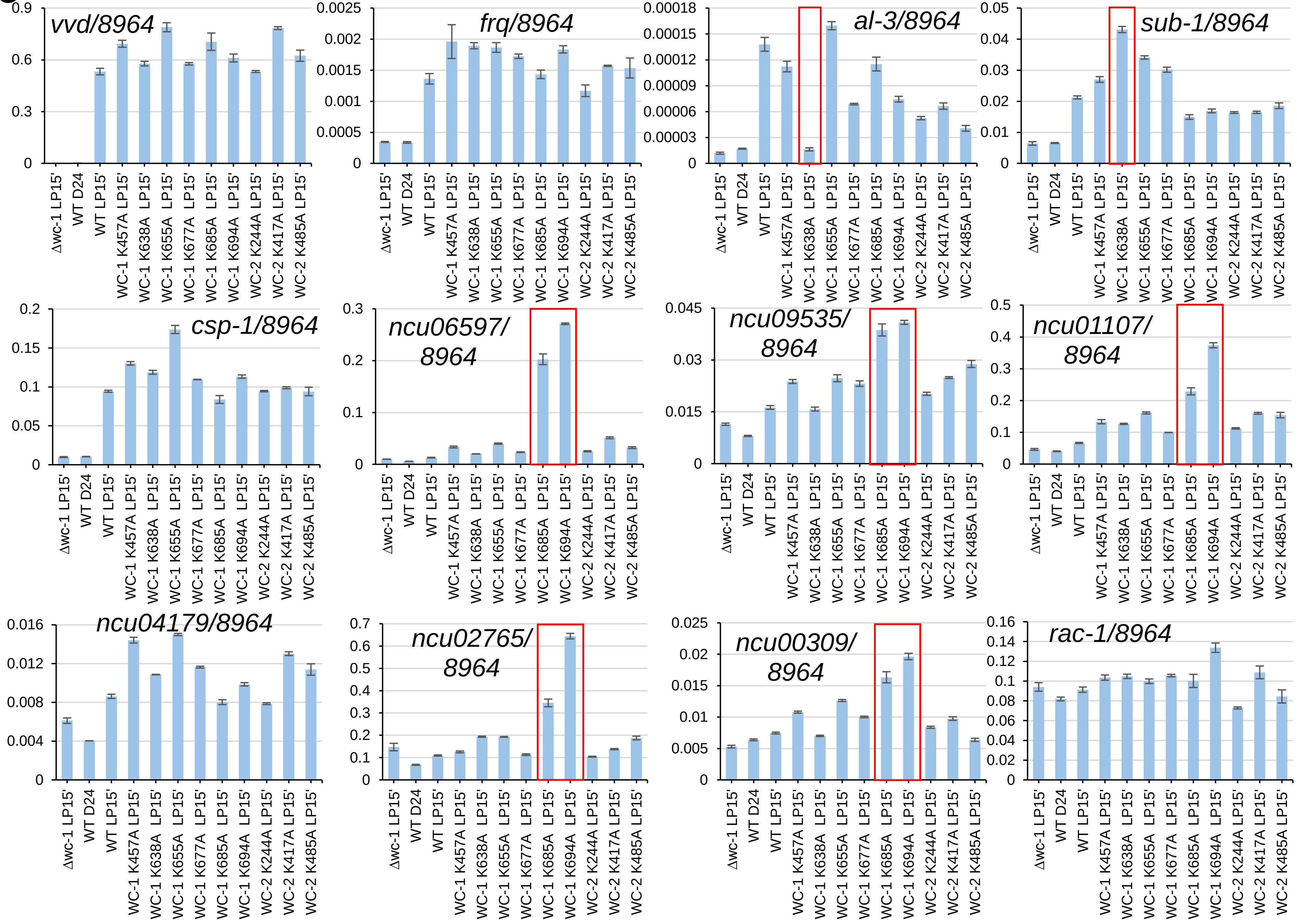
