## Supplementary material for "Acetylation of WCC is dispensable for the core circadian clock but differentially regulates acute light responses in *Neurospora*": suppl. table S2

| sequence | ref | descripti | charge |
|---|---|---|---|
| R.CTSWQYELR!.Q | sp\|Q01371\|WC1 | White col | 2 |
| K.DLELNGGIGDSEIWTK!.V | sp\|Q01371\|WC1 | White col | 2 |
| K.TIAEGQEIQQSLINYR!.K | sp\|Q01371\|WC1 | White col | 2 |
| R.FLQAPDGNVEAGTK!.R | sp\|Q01371\|WC1 | White col | 2 |
| R.FLQAPDGNVEAGTK!R.E | sp\|Q01371\|WC1 | White col | 3 |
| K.GLFLYLSPACK!.K | sp\|Q01371\|WC1 | White col | 2 |
| K.QLEPADGQTLGVDDVSTLLQQCNSK!.G | sp\|Q01371\|WC1 | White col | 3 |
| R.KDLELNGGIGDSEIWTK!.V | sp\|Q01371\|WC1 | White col | 3 |
| K.R!EFVENNAVYTLKK.T | sp\|Q01371\|WC1 | White col | 3 |
| K.SGSSDSTGAM*VSAR!.S | sp\|Q01371\|WC1 | White col | 2 |
| K.VSTSGM*FLFVSSNVR!.S | sp\|Q01371\|WC1 | White col | 2 |

| match | tp | xcorr_ma | dcn_max | ppm_min | methyl | non-methyl |
|---|---|---|---|---|---|---|
| 1 | 6 | 2.166 | 0.8508 | 0.170495 | http://or | http://orion.dartmouth.edu/ms20/spar |
| 1 | 3 | 3.06 | 0.9203 | 1.636827 | http://or | http://orion.dartmouth.edu/ms20/spar |
| 1 | 3 | 3.181 | 0.8236 | 0.742556 | http://or | http://orion.dartmouth.edu/ms20/spar |
| 1 | 7 | 2.296 | 0.9226 | 0.259622 | http://or | http://orion.dartmouth.edu/ms20/spar |
| 1 | 6 | 3.289 | 0.9696 | 0.885592 | http://or | http://orion.dartmouth.edu/ms20/spar |
| 1 | 5 | 2.608 | 0.8981 | 0.046964 | http://or | http://orion.dartmouth.edu/ms20/spar |
| 1 | 2 | 3.281 | 0.9828 | 0.536881 | http://or | http://orion.dartmouth.edu/ms20/spar |
| 1 | 1 | 2.237 | 0.8605 | 1.309108 | http://or | http://orion.dartmouth.edu/ms20/spar |
| 1 | 1 | 2.075 | 0.8916 | 1.003436 | http://or | http://orion.dartmouth.edu/ms20/spar |
| 1 | 1 | 3.321 | 0.9509 | 1.7025 | http://or | http://orion.dartmouth.edu/ms20/spar |
| 1 | 1 | 1.956 | 0.8952 | 0.86147 | http://or | http://orion.dartmouth.edu/ms20/spar |

row?psm=1239075141
row?psm=1239073728
row?psm=1238884565
row?psm=1238884082
row?psm=1238883932
row?psm=1238882568
row?psm=1238856973
row?psm=1238856970
row?psm=1238855474
row?psm=1238736537
row?psm=1238884650
