## Supplementary material for "Acetylation of WCC is dispensable for the core circadian clock but differentially regulates acute light responses in *Neurospora*": suppl. table S1

| search | sample | fraction | scan | charge | precursor | precursor_ | precursor | ions_matc |
|---|---|---|---|---|---|---|---|---|
| 82854 | 19021 | | 5977 | 4 | 5972 | 328008 | 552.7857 | 12 |
| 82856 | 29096 | | 5677 | 2 | 5674 | 2611700 | 476.2304 | 7 |
| 82869 | 19087 | | 742 | 3 | 734 | 842304 | 516.5868 | 10 |
| 82878 | 30006 | | 5587 | 3 | 5583 | 379556 | 516.5868 | 6 |
| 82856 | 29096 | | 4039 | 2 | 4037 | 502805 | 356.7031 | 8 |
| 82892 | 20572 | | 7386 | 3 | 7375 | 5012460 | 678.3611 | 22 |
| 82890 | 34461 | | 5389 | 3 | 5386 | 712630 | 678.3606 | 25 |
| 82887 | 20570 | | 6681 | 3 | 6675 | 656301 | 678.3609 | 16 |
| 82864 | 19029 | | 3830 | 2 | 3826 | 535264 | 1017.038 | 19 |
| 82890 | 34461 | | 5560 | 3 | 5555 | 489094 | 678.3599 | 18 |
| 82899 | 20576 | | 6057 | 3 | 6051 | 603379 | 678.361 | 14 |
| 82882 | 20568 | | 8237 | 3 | 8232 | 530900 | 678.3608 | 10 |
| 82864 | 19029 | | 3827 | 3 | 3825 | 597394 | 678.361 | 9 |
| 82896 | 20574 | | 5662 | 3 | 5655 | 614040 | 721.0596 | 21 |
| 82892 | 20572 | | 6173 | 3 | 6166 | 539694 | 721.0591 | 12 |
| 82899 | 20576 | | 5051 | 3 | 5043 | 412808 | 721.0596 | 9 |
| 82896 | 20574 | | 6247 | 3 | 6236 | 2916610 | 651.0189 | 22 |
| 82890 | 34461 | | 5160 | 3 | 5157 | 714500 | 651.0189 | 18 |
| 82842 | 18987 | | 4074 | 3 | 4070 | 835856 | 651.0188 | 15 |
| 82899 | 20576 | | 5523 | 4 | 5521 | 771326 | 488.5157 | 15 |
| 82892 | 20572 | | 6748 | 3 | 6731 | 4753240 | 651.0189 | 15 |
| 82862 | 19027 | | 2828 | 3 | 2824 | 484351 | 651.0186 | 13 |
| 82844 | 19002 | | 3068 | 3 | 3062 | 694249 | 651.019 | 10 |
| 82896 | 20574 | | 6266 | 4 | 6257 | 917904 | 488.5154 | 8 |
| 82882 | 20568 | | 7510 | 4 | 7494 | 938138 | 488.5155 | 7 |
| 82832 | 29996 | | 16037 | 3 | 16030 | 368158 | 651.0184 | 8 |
| 82866 | 29996 | | 16037 | 3 | 16030 | 368158 | 651.0184 | 8 |
| 82902 | 29996 | | 16037 | 3 | 16030 | 368158 | 651.0184 | 8 |
| 82841 | 29996 | | 16037 | 3 | 16030 | 368158 | 651.0184 | 8 |
| 82864 | 19029 | | 3443 | 3 | 3438 | 479625 | 651.0185 | 8 |
| 82899 | 20576 | | 5509 | 3 | 5503 | 982085 | 651.0192 | 10 |
| 82887 | 20570 | | 6054 | 4 | 6038 | 1062650 | 488.5156 | 8 |
| 82859 | 19025 | | 3333 | 2 | 3331 | 548472 | 500.8027 | 11 |
| 82844 | 19002 | | 2669 | 2 | 2667 | 572425 | 500.8027 | 10 |
| 82892 | 20572 | | 5745 | 2 | 5738 | 1286360 | 500.8027 | 10 |
| 82896 | 20574 | | 5230 | 2 | 5227 | 1648070 | 500.8027 | 10 |
| 82887 | 20570 | | 5271 | 2 | 5265 | 1759390 | 500.8027 | 10 |
| 82842 | 18987 | | 3760 | 2 | 3758 | 507744 | 500.8026 | 10 |
| 82864 | 19029 | | 2977 | 2 | 2975 | 593754 | 500.8027 | 10 |
| 82899 | 20576 | | 4676 | 2 | 4672 | 960044 | 500.8026 | 9 |
| 82896 | 20574 | | 4968 | 2 | 4957 | 868066 | 500.8021 | 6 |
| 82896 | 20574 | | 7964 | 3 | 7957 | 1244500 | 639.6589 | 18 |
| 82892 | 20572 | | 8489 | 3 | 8473 | 583196 | 639.6589 | 14 |
| 82896 | 20574 | | 7966 | 2 | 7957 | 865772 | 958.9851 | 7 |
| 82882 | 20568 | | 9405 | 3 | 9399 | 639981 | 639.659 | 4 |
| 82842 | 18987 | | 4576 | 3 | 4573 | 475682 | 791.3708 | 20 |
| 82864 | 19029 | | 4161 | 3 | 4159 | 515605 | 791.3709 | 24 |

| | | | | | | | |
|---|---|---|---|---|---|---|---|
| 82892 | 20572 | 8118 | 4 | 8114 | 336402 | 593.7798 | 18 |
| 82899 | 20576 | 6677 | 4 | 6672 | 680057 | 593.7796 | 18 |
| 82896 | 20574 | 7619 | 4 | 7608 | 625360 | 593.7797 | 18 |
| 82896 | 20574 | 7613 | 3 | 7605 | 761752 | 791.3707 | 18 |
| 82892 | 20572 | 8105 | 3 | 8101 | 349444 | 791.3709 | 16 |
| 82844 | 19002 | 3642 | 3 | 3639 | 429558 | 791.3709 | 13 |
| 82899 | 20576 | 6667 | 3 | 6663 | 426476 | 791.3708 | 8 |
| 82859 | 19025 | 2122 | 3 | 2119 | 529791 | 389.2366 | 19 |
| 82892 | 20572 | 3062 | 3 | 3045 | 3474050 | 389.2367 | 16 |

| ions_made | rt | ppm | xcorr | dcn | psc | enzymati | ref | duplicate |
|---|---|---|---|---|---|---|---|---|
| 114 | 2120.71 | 0.662319 | 2.377 | 0.7981 | 5 | | sp\|P78714\|WC2_NEUC | |
| 12 | 1551.86 | 0.575162 | 1.528 | 0.6859 | 3 | | sp\|P78714\|WC2_NEUC | |
| 48 | 863.35 | 0.636965 | 1.575 | 0.4337 | 4 | | sp\|P78714\|WC2_NEUC | |
| 48 | 1691.45 | 0.54005 | 1.123 | 0.3188 | 4 | | sp\|P78714\|WC2_NEUC | |
| 12 | 1322.86 | 0.208086 | 1.634 | 0.7001 | 3 | | sp\|P78714\|WC2_NEUC | |
| 64 | 2008.33 | 1.299446 | 3.907 | 0.8329 | 3 | | sp\|Q01371\|WC1_NEUC | |
| 64 | 2131.39 | 0.546888 | 3.898 | 0.8486 | 3 | | sp\|Q01371\|WC1_NEUC | |
| 64 | 2019.65 | 0.989569 | 3.411 | 0.8519 | 3 | | sp\|Q01371\|WC1_NEUC | |
| 32 | 2060.85 | 1.469819 | 3.382 | 0.7439 | 3 | | sp\|Q01371\|WC1_NEUC | |
| 64 | 2167.83 | -0.45652 | 3.182 | 0.9029 | 3 | | sp\|Q01371\|WC1_NEUC | |
| 64 | 2015.15 | 1.151886 | 2.853 | 0.8829 | 3 | | sp\|Q01371\|WC1_NEUC | |
| 64 | 2026.51 | 0.812497 | 2.485 | 0.7996 | 3 | | sp\|Q01371\|WC1_NEUC | |
| 64 | 2060.31 | 1.13713 | 2.008 | 0.7535 | 3 | | sp\|Q01371\|WC1_NEUC | |
| 68 | 1840.38 | 1.387615 | 3.904 | 0.9813 | 4 | | sp\|Q01371\|WC1_NEUC | |
| 68 | 1851.62 | 0.693544 | 2.294 | 0.9891 | 4 | | sp\|Q01371\|WC1_NEUC | |
| 68 | 1853.58 | 1.484785 | 1.765 | 0.9813 | 4 | | sp\|Q01371\|WC1_NEUC | |
| 60 | 1924.12 | 1.26592 | 2.711 | 0.8746 | 4 | | sp\|Q01371\|WC1_NEUC | |
| 60 | 2073.85 | 1.327426 | 2.697 | 0.8899 | 4 | | sp\|Q01371\|WC1_NEUC | |
| 60 | 2183.35 | 1.142909 | 2.669 | 0.7171 | 4 | | sp\|Q01371\|WC1_NEUC | |
| 90 | 1929.6 | 0.647593 | 2.415 | 0.7379 | 4 | | sp\|Q01371\|WC1_NEUC | |
| 60 | 1930.3 | 1.235167 | 2.204 | 0.8367 | 4 | | sp\|Q01371\|WC1_NEUC | |
| 60 | 1960.42 | 0.866133 | 2.095 | 0.5446 | 4 | | sp\|Q01371\|WC1_NEUC | |
| 60 | 1975.24 | 1.496566 | 2.033 | 0.6572 | 4 | | sp\|Q01371\|WC1_NEUC | |
| 90 | 1925.98 | 0.17605 | 1.971 | 0.8564 | 4 | | sp\|Q01371\|WC1_NEUC | |
| 90 | 1943 | 0.340065 | 1.713 | 0.7963 | 4 | | sp\|Q01371\|WC1_NEUC | |
| 60 | 3105.28 | 0.573981 | 1.651 | 0.6905 | 4 | | sp\|Q01371\|WC1_NEUC | |
| 60 | 3105.28 | 0.573981 | 1.651 | 0.6905 | 4 | | sp\|Q01371\|WC1_NEUC | |
| 60 | 3105.28 | 0.573981 | 1.651 | 0.8383 | 4 | | sp\|Q01371\|WC1_NEUC | |
| 60 | 3105.28 | 0.573981 | 1.651 | 0.8383 | 4 | | sp\|Q01371\|WC1_NEUC | |
| 60 | 1952.76 | 0.681616 | 1.561 | 0.6067 | 4 | | sp\|Q01371\|WC1_NEUC | |
| 60 | 1927.28 | 1.834847 | 1.461 | 0.7734 | 4 | | sp\|Q01371\|WC1_NEUC | |
| 90 | 1919.75 | 0.463076 | 1.336 | 0.8159 | 4 | | sp\|Q01371\|WC1_NEUC | |
| 14 | 1808.02 | 0.848733 | 2.236 | 0.7093 | 3 | | sp\|Q01371\|WC1_NEUC | |
| 14 | 1824.6 | 0.868721 | 2.214 | 0.7484 | 3 | | sp\|Q01371\|WC1_NEUC | |
| 14 | 1790.95 | 1.008638 | 2.198 | 0.7812 | 3 | | sp\|Q01371\|WC1_NEUC | |
| 14 | 1780.27 | 0.98865 | 2.179 | 0.6829 | 3 | | sp\|Q01371\|WC1_NEUC | |
| 14 | 1807.21 | 0.888709 | 2.166 | 0.8241 | 3 | | sp\|Q01371\|WC1_NEUC | |
| 14 | 2083.39 | 0.808757 | 2.152 | 0.7165 | 3 | | sp\|Q01371\|WC1_NEUC | |
| 14 | 1804.27 | 0.98865 | 2.14 | 0.7294 | 3 | | sp\|Q01371\|WC1_NEUC | |
| 14 | 1785.51 | 0.788769 | 2.019 | 0.7568 | 3 | | sp\|Q01371\|WC1_NEUC | |
| 14 | 1745.31 | -0.35055 | 1.221 | 0.5315 | 3 | | sp\|Q01371\|WC1_NEUC | |
| 64 | 2151.17 | 1.1674 | 3.455 | 0.8515 | 3 | | sp\|Q01371\|WC1_NEUC | |
| 64 | 2165.16 | 1.151751 | 3.197 | 0.8586 | 3 | | sp\|Q01371\|WC1_NEUC | |
| 32 | 2151.51 | 1.546323 | 1.822 | 0.933 | 3 | | sp\|Q01371\|WC1_NEUC | |
| 64 | 2171.34 | 1.339548 | 1.33 | 0.912 | 3 | | sp\|Q01371\|WC1_NEUC | |
| 76 | 2317.31 | 1.549626 | 4.073 | 0.7793 | 4 | | sp\|Q01371\|WC1_NEUC | |
| 76 | 2157.52 | 1.688744 | 3.974 | 0.7967 | 4 | | sp\|Q01371\|WC1_NEUC | |

| | | | | | | |
|---|---|---|---|---|---|---|
| 114 | 2106.01 | 1.302427 | 3.456 | 0.8892 | 4 | sp\|Q01371\|WC1_NEU( |
| 114 | 2130.55 | 1.032623 | 3.455 | 0.9045 | 4 | sp\|Q01371\|WC1_NEU( |
| 114 | 2107.91 | 1.167525 | 3.427 | 0.9212 | 4 | sp\|Q01371\|WC1_NEU( |
| 76 | 2107.25 | 1.347274 | 2.887 | 0.974 | 4 | sp\|Q01371\|WC1_NEU( |
| 76 | 2103.45 | 1.688744 | 2.737 | 0.9678 | 4 | sp\|Q01371\|WC1_NEU( |
| 76 | 2175.25 | 1.688744 | 2.205 | 0.7138 | 4 | sp\|Q01371\|WC1_NEU( |
| 76 | 2128.92 | 1.499038 | 1.834 | 0.9831 | 4 | sp\|Q01371\|WC1_NEU( |
| 32 | 1432.27 | -0.07304 | 3.027 | 0.6944 | 5 | sp\|Q01371\|WC1_NEU( |
| 32 | 1402.79 | 0.107112 | 2.787 | 0.7234 | 5 | sp\|Q01371\|WC1_NEU( |

| decoy | sequence | DOUBLE CHECK # site |
|---|---|---|
| 0 | K.K~DGNWTIFETVGHAHIAAAK.F | K244 |
| 0 | L.K~YQEGER.S | K417 |
| 0 | L.K~YQEGERSHGITT.G | K417 |
| 0 | L.K~YQEGERSHGITT.G | K417 |
| 0 | R.K~GPSGPK.T | K481 |
| 0 | K.K~TIAEGQEIQQSLINYR.K | K457 |
| 0 | K.K~TIAEGQEIQQSLINYR.K | K457 |
| 0 | K.K~TIAEGQEIQQSLINYR.K | K457 |
| 0 | K.K~TIAEGQEIQQSLINYR.K | K457 |
| 0 | K.K~TIAEGQEIQQSLINYR.K | K457 |
| 0 | K.K~TIAEGQEIQQSLINYR.K | K457 |
| 0 | K.K~TIAEGQEIQQSLINYR.K | K457 |
| 0 | K.K~TIAEGQEIQQSLINYR.K | K457 |
| 0 | K.K~TIAEGQEIQQSLINYRK.G | K457 |
| 0 | K.K~TIAEGQEIQQSLINYRK.G | K457 |
| 0 | K.K~TIAEGQEIQQSLINYRK.G | K457 |
| 0 | R.ELK~EAQQHTPVNIVFR.I | K638 |
| 0 | R.ELK~EAQQHTPVNIVFR.I | K638 |
| 0 | R.ELK~EAQQHTPVNIVFR.I | K638 |
| 0 | R.ELK~EAQQHTPVNIVFR.I | K638 |
| 0 | R.ELK~EAQQHTPVNIVFR.I | K638 |
| 0 | R.ELK~EAQQHTPVNIVFR.I | K638 |
| 0 | R.ELK~EAQQHTPVNIVFR.I | K638 |
| 0 | R.ELK~EAQQHTPVNIVFR.I | K638 |
| 0 | R.ELK~EAQQHTPVNIVFR.I | K638 |
| 0 | R.ELK~EAQQHTPVNIVFR.I | K638 |
| 0 | R.ELK~EAQQHTPVNIVFR.I | K638 |
| 0 | R.ELK~EAQQHTPVNIVFR.I | K638 |
| 0 | R.ELK~EAQQHTPVNIVFR.I | K638 |
| 0 | R.ELK~EAQQHTPVNIVFR.I | K638 |
| 0 | R.ELK~EAQQHTPVNIVFR.I | K638 |
| 0 | R.K~CIILVGR.K | K677 |
| 0 | R.K~CIILVGR.K | K677 |
| 0 | R.K~CIILVGR.K | K677 |
| 0 | R.K~CIILVGR.K | K677 |
| 0 | R.K~CIILVGR.K | K677 |
| 0 | R.K~CIILVGR.K | K677 |
| 0 | R.K~CIILVGR.K | K677 |
| 0 | R.K~CIILVGR.K | K677 |
| 0 | R.K~CIILVGR.K | K677 |
| 0 | R.K~DLELNGGIGDSEIWTK.V | K694 |
| 0 | R.K~DLELNGGIGDSEIWTK.V | K694 |
| 0 | R.K~DLELNGGIGDSEIWTK.V | K694 |
| 0 | R.K~DLELNGGIGDSEIWTK.V | K694 |
| 0 | R.K~NSGYTWFESHGTLFNEQGK.G | K655 |
| 0 | R.K~NSGYTWFESHGTLFNEQGK.G | K655 |

| | | |
|---|---|---|
| 0 | R.K~NSGYTWFESHGTLFNEQGK.G | K655 |
| 0 | R.K~NSGYTWFESHGTLFNEQGK.G | K655 |
| 0 | R.K~NSGYTWFESHGTLFNEQGK.G | K655 |
| 0 | R.K~NSGYTWFESHGTLFNEQGK.G | K655 |
| 0 | R.K~NSGYTWFESHGTLFNEQGK.G | K655 |
| 0 | R.K~NSGYTWFESHGTLFNEQGK.G | K655 |
| 0 | R.K~NSGYTWFESHGTLFNEQGK.G | K655 |
| 0 | R.K~RPVFALHR.K | K685 |
| 0 | R.K~RPVFALHR.K | K685 |

| | |
|---|---|
| http://orion.dartmou | MAYBE |
| http://orion.dartmou | MAYBE |
| http://orion.dartmou | MAYBE |
| | MAYBE |
| http://orion.dartmou | MAYBE |

http://orion.dartmouth.edu/ms20/sparrow?psm=1238195307

http://orion.dartmouth.edu/ms20/sparrow?psm=1238208029

http://orion.dartmouth.edu/ms20/sparrow?psm=1237673355

http://orion.dartmouth.edu/ms20/sparrow?psm=1237842892

http://orion.dartmouth.edu/ms20/sparrow?psm=1238208905

http://orion.dartmouth.edu/ms20/sparrow?psm=1237900298

http://orion.dartmouth.edu/ms20/sparrow?psm=1237958198

http://orion.dartmouth.edu/ms20/sparrow?psm=1237842447
